# ER-to-Golgi trafficking of procollagen in the absence of large carriers

**DOI:** 10.1101/339804

**Authors:** Janine McCaughey, Nicola L. Stevenson, Stephen Cross, David J. Stephens

## Abstract

Secretion and assembly of collagen is fundamental to the function of the extracellular matrix. Defects in the assembly of a collagen matrix lead to pathologies including fibrosis and osteogenesis imperfecta. Owing to the size of fibril-forming procollagen molecules it is assumed that they are transported from the endoplasmic reticulum to the Golgi in specialised large COPII-dependent carriers. Here, analysing endogenous procollagen and a new engineered GFP-tagged form, we show that transport to the Golgi occurs in the absence of large carriers. Large GFP-positive structures are observed occasionally but these are non-dynamic, are not COPII-positive, and label with markers of the ER. We propose a “short-loop” model of COPII-dependent ER-to-Golgi traffic that, while consistent with models of ERGIC-dependent expansion of COPII carriers, does not invoke long-range trafficking of large vesicular structures. Our findings provide an important insight into the process of procollagen trafficking and reveal a short-loop pathway from the ER to the Golgi, without the use of large carriers.

**Summary:** Trafficking of procollagen is essential for normal cell function. Here, imaging of GFP-tagged type I procollagen reveals that it is transported from the endoplasmic reticulum to the Golgi, without the use of large carriers.

## Introduction

Collagen is the most abundant protein in the body. Fibrillar type I collagen plays a key role in bone, skin and tendon formation, providing tissues with the necessary structural support. Altered collagen secretion, processing, and assembly is linked to diseases including osteogenesis imperfecta (OI), fibrosis, chondrodysplasia, Ehlers-Danlos syndrome and many more (Forlino and Marini, 2016; Jobling et al., 2014). Type I collagen assembles from two type I α1 chains together with one type I α2 chain to form trimeric procollagen in the endoplasmic reticulum (ER) (Canty and Kadler, 2005; Goldberg et al., 1972), with the α-helix from each chain forming a rigid 300 nm triple helix structure (Bachinger et al., 1982; Lightfoot et al., 1992). During procollagen biosynthesis proline hydroxylation stabilises the triple helical conformation (Blanck and Peterkofsky, 1975; Jimenez et al., 1973). This process requires the presence of ascorbic acid, which acts as a co-factor for prolyl-4-hydroxylase (Mussini et al., 1967). The collagen-specific chaperone *heat shock protein 47* (Hsp47) (Ito and Nagata, 2017; Satoh et al., 1996) is also required.

To be secreted efficiently, procollagen I must traffic from the ER to the Golgi via the ER-Golgi intermediate compartment (ERGIC) (Malhotra and Erlmann, 2015; Malhotra et al., 2015; Satoh et al., 1996). Conventional ER-to-Golgi transport is facilitated by coat protein complex type II (COPII) vesicles with a size of 60 - 90 nm in diameter. These vesicles are thus significantly smaller than the 300 nm length of procollagen. Nonetheless, COPII vesicles are essential for efficient collagen trafficking in cells (Stephens and Pepperkok, 2002; Townley et al., 2008; Townley et al., 2012) and in animal models, since perturbation of, or mutations in, key COPII components including Sec24D (Garbes et al., 2015; Moosa et al., 2015; Sarmah et al., 2010), Sec23A (Boyadjiev et al., 2006; Lang et al., 2006), and Sec13 (Schmidt et al., 2013; Townley et al., 2008; Townley et al., 2012) cause defects in collagen secretion. To accommodate all of these data the prevailing hypothesis for the mechanism of procollagen secretion proposes the formation of large COPII carriers (Gorur et al., 2017; Malhotra and Erlmann, 2015; McGourty et al., 2016; Nogueira et al., 2014; Saito et al., 2009; Saito and Katada, 2015; Santos et al., 2015; Venditti et al., 2012). Formation of these carriers is said to be facilitated by the ER transmembrane proteins TANGO1 (*transport and Golgi organisation 1*, also called Mia3) and cTAGE5 (a TANGO1-related protein) that form a dimer and localises to ER exit sites (ERES) in mammals (Saito et al., 2011). TANGO1 is considered to act as a tether to the ERGIC to expand the nascent procollagen-containing carrier during its formation.

Several publications have shown large structures reported to be ER-to-Golgi carriers of procollagen (Gorur et al., 2017; Jin et al., 2012; McGourty et al., 2016). These large structures are generally few in number and seen in systems where the ubiquitin ligase Cullin3 (CUL3) adaptor Kelch-like protein 12 (KLHL12) is overexpressed (Jin et al., 2012). CUL3 facilitates mono-ubiquitylation of Sec31, stalling the outer complex formation, leading to a delayed scission and enlargement of COPII vesicles (Jin et al., 2012). Subsequent work defined a mechanism for KLHL12-mediated ubiquitylation of Sec31A, with PEF1 and ALG2 shown to be subunits of the CUL3-KLHL12 ubiquitin ligase (McGourty et al., 2016). Consistent with the previous finding, the large structures labelled for Sec31A, PEF1, and ALG-2 and, more significantly, with antibodies against procollagen (McGourty et al., 2016). Similar large, procollagen- and Sec31A-positive structures have been shown in KI6 cells that overexpress both type I procollagen and FLAG-KLHL12 (Gorur et al., 2017). Apparent encapsulation of procollagen into large Sec31A structures was shown by correlative light electron microscopy (Gorur et al., 2017). Interestingly, smaller Sec31A puncta remained devoid of procollagen labelling. Short-range movement of these carriers, using live cell imaging of procollagen-GFP, did not appear to be Golgi-directed and only occurred over distances of a few microns. Further support for this model came from experiments showing COPII-dependent packaging of procollagen *in vitro* into large structures (Gorur et al., 2017; Yuan et al., 2017).

Pathways for transport of procollagen from the ER to the Golgi have to date relied extensively on a COL1A1-GFP construct with a C-terminal GFP-tag (Stephens and Pepperkok, 2002). Expression of this construct reveals the presence of small ascorbate-dependent, highly mobile long-range transport carriers containing procollagen-GFP. While generally effective there are caveats to its use. Many cells do not traffic this fusion protein effectively. This compromised ability to assemble and traffic is most likely because the folding of the procollagen trimer proceeds from the C-terminus (Bourhis et al., 2012; Kadler et al., 1990).

Our aim was to test the prevailing hypothesis that large vesicular carriers mediate procollagen secretion. To achieve this, we generated a newly designed procollagen construct incorporating some key features to better control and define secretory cargo export from the ER. Surprisingly, live cell imaging using this construct revealed that procollagen traffics from the ER to the Golgi without using large vesicular carriers. Instead the Golgi gradually fills with procollagen upon ascorbate addition, while the ER empties, suggesting the existence of a short trafficking loop connecting the ER and Golgi. Combined with work on endogenous procollagen and a variety of cell types we propose a new mode of procollagen trafficking via a short-loop pathway that facilitates local transfer of procollagen from the juxtanuclear ER to the Golgi.

## Results

### An ascorbate- and biotin-controllable procollagen reporter: mGFP-SBP-COL1A1

To visualise ER-to-Golgi trafficking of procollagen for study, we engineered monomeric GFP (mGFP) and a streptavidin-binding peptide (SBP)-tag into procollagen 1α1. The tag was introduced upstream of the naturally occurring N-terminal proteolytic cleavage site (located after the N-terminal pro-peptide) to generate mGFP-SBP-COL1A1 (abbreviated here to GFP-COL, Fig. 1A). The inclusion of the SBP-tag allows synchronisation of trafficking using the Retention Using Selective Hooks (RUSH) system to control cargo exit from the ER using biotin (Boncompain et al., 2012). When expressed transiently in IMR-90 human lung fibroblasts (Fig. 1Bi), and in human telomerase immortalised retinal pigment epithelial cells (hTERT-RPE-1, Fig. 1Bii), GFP-COL colocalises with Hsp47 in the ER, but not with the *cis*-Golgi marker GM130 at steady state (Fig. 1Ci). Following the addition of ascorbate, GFP-COL became enriched in the Golgi area (Fig. 1Cii - iii).

**Figure 1:**
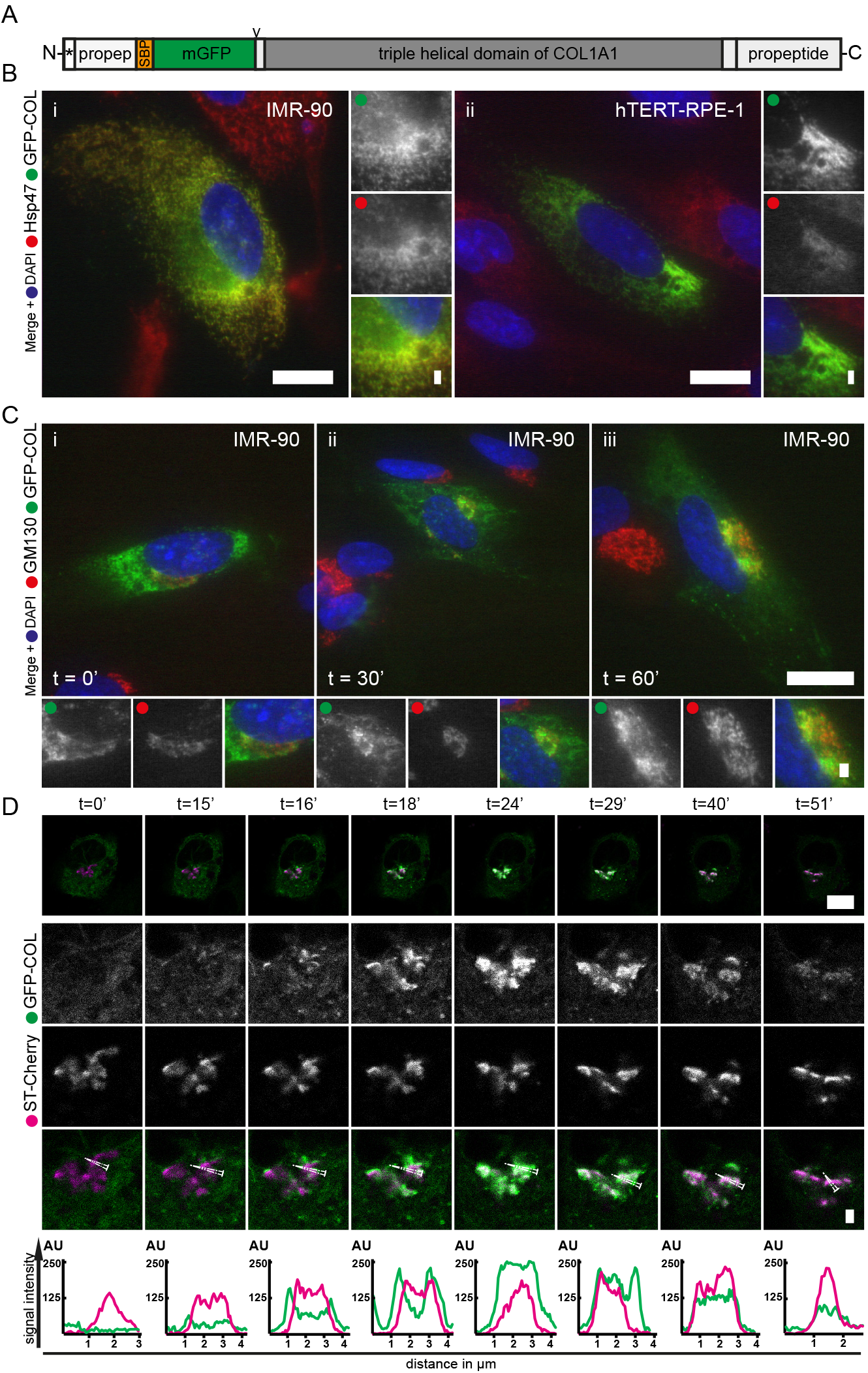
Controllable ER-to-Golgi traffcking of GFP-COL in the absence of large carriers. The GFP-COL fusion construct is shown in A. It consists of a synthetic pro-sequence of human COL1A1 followed by a streptavi-din-binding peptide (SBP), monomeric green fluorescent protein (mGFP), the N-propeptide cleavage site (indicated by the rrowhead) within the non-helical region and the triple-helical domain with the C-terminal nonhelical region followed by the C-propeptide of human COL1A1. The signal sequence is indicated by *. Widefield microscopy of cells transiently expressing GFP-COL (A) is shown in B and C. 16 hours after transfection, GFP-COL (green) colocalises with Hsp47 (red) in the ER in IMR-90 cells (Bi) and RPE-1 (Bii). Large images show whole cells, while the small panels show the enlargements with the corresponding channels in greyscale and the merge image including nuclear DAPI staining (blue). C: Shows same image set-up as in B with GFP-COL (green) and cis-Golgi marker GM130 (red) labelling in transiently GFP-COL expressing IMR-90 cells. Timepoints indicate length of incubation in the presence of 50 μg.ml-1 ascorbate prior to fixation. A concentration of GFP-COL in the Golgi can be observed after both 30 (Cii) and 60 mins (Ciii) in presence with ascorbate. Number of cells imaged n ≥ 10. D: Confocal images from live cell imaging of RPE-1 stably expressing GFP-COL (green; GFP-COL-RPE) co-transfected with the trans-Golgi marker ST-Cherry (magenta; utilising the RUSH system). Acquisition at 1 frame every 30 seconds. Image layout as in C, followed by line line with a 5-pixel width drawn through the Golgi in the enlarged overlays. X-axis shows the distance in μm. Image stills are derived from Video 1. Timepoints indicate mins of incubation in presence of 500 μg.ml-1 ascorbate and 400 μM biotin (asc/biotin). At timepoint 0 (t = 0 mins) GFP-COL and ST-Cherry show distinct localisation. Over time an accumulation of GFP-COL around the Golgi marker occurs and increases until t = 18 mins, with subsequent filling of the trans-Golgi (t = 24 mins). Later time points show the decrease in intensity of GFP-COL overall and in the Golgi area (t = 29, 40 and 51 mins), implying functional transport to the Golgi, followed by secretion from the cell. For each set of live imaging experiment 3 cells from the same dish were imaged simultaneously. A total of n = 4 sets was acquired. Scale bars = 10 μm and 1 μm (in enlargements).

As expected, transient transfection of GFP-COL-positive cells led to highly variable expression levels. To reduce the variability a RPE-1 cell line was created that stably expresses GFP-COL (GFP-COL-RPE). The population was sorted into four equally distributed populations with different expression levels according to signal intensity of GFP and the 25% of cells expressing the lowest level was chosen for future experiments. For further analysis of transport to the Golgi GFP-COL-RPE cells were co-transfected with a bicistronic construct encoding an ER-hook (KDEL-tagged streptavidin to retain the SBP fusion protein in the ER), and a separate *trans*-Golgi marker that expressed the minimal Golgi targeting region of sialyltransferase fused to mCherry (ST-Cherry). The co-expression of the ER-hook here enabled the retention of SBP-tagged GFP-COL in the ER lumen until addition of biotin to trigger the dissociation of SBP from the streptavidin-bound KDEL “hook”.

Live cell imaging of synchronised procollagen transport using confocal microscopy was then used to monitor the dynamics of ER-to-Golgi transport. Prior to addition of ascorbate and biotin (asc/biotin) GFP-COL and the Golgi marker show no significant colocalization (Fig. 1D, t=0). After addition of asc/biotin GFP-COL structures of high signal intensity appeared near the Golgi, as well as in the cell periphery (Fig. 1D, t=15 and Video 1). The signal intensity of GFP-COL in the ER decreased gradually over time while the accumulation of GFP-COL in and around the *trans*-Golgi further increased (Video 1). After filling of the Golgi an overall decrease of GFP-COL signal intensity occurred over time. Line scans in Fig. 1D show clearly this accumulation of GFP-COL around the ST-Cherry labelled Golgi compartments (t=16 mins), before GFP-COL fills the *trans*-Golgi (here at 24 mins asc/biotin), and subsequently exits (t=40 mins and 51 mins in this example). These experiments are consistent with ascorbate-dependent transport of GFP-COL to the Golgi, followed by its secretion from the cell. Supplemental Figure S1A (derived from Video 2) shows that the addition of ascorbate alone is insufficient to release GFP-COL when co-expressed with a hook (in this case a KDEL-tagged streptavidin as ER-hook and separate *cis*-Golgi marker mannosidase II tagged with mScarlet-i, a red fluorescent protein (Bindels et al., 2017); MannII-mSc). Supplemental Figure S1 shows ascorbate- and biotin-dependent trafficking in hTERT-BJ5Ta human fibroblasts (Fig. S1B and Video 3). Ascorbate-dependent trafficking in IMR-90 human fibroblasts is shown in Fig. S1C and Video 4.

Surprisingly, in the above experiments we failed to observe the formation of any large, dynamic carriers prior to procollagen reaching the Golgi. Imaging at a higher frame rate (Fig. S2 and Video 5) also failed to reveal any large puncta that translocate towards the Golgi. In some experiments (more noticeably in transiently transfected cells) we did detect a very small number (typically 1 per cell) of punctate structures that translocate to the Golgi along curvilinear tracks, consistent with previously described vesicular tubular structures (Stephens and Pepperkok, 2002). An example of this is Fig. S1B and Video 3, where a GFP-positive structure with a maximum estimated size of 535 nm translocates to the Golgi over the nucleus at ^~^17 mins post-asc/biotin. Similarly, Fig. S2 and Video 5 show one such structure tracking towards to the Golgi over the nucleus (highlighted by arrowheads 10’03’’-11’00’’). Collectively these data suggest ER-to-Golgi transport in the absence of large carriers. We therefore sought to define this in more detail.

### ER-to-Golgi trafficking of GFP-COL occurs without the use of large carriers

In approximately 50% of all live cells, we did observe between one and four large, round GFP-COL structures (calculated diameter >1 μm) that were similar in appearance to those previously reported. For example, Fig. 2A and 2B show GFP-COL distributed throughout the ER and in four larger (1 - 2 μm diameter), circular, GFP-positive structures with elevated signal intensity compared to the ER background (Fig. 2B, circles). However, these structures were relatively static and did not change in size or intensity despite transport of GFP-COL to the Golgi (Fig. 2A-D and Videos 6 and 7). They also did not appear to be the source or destination of the few more dynamic punctate structures observed (e.g. highlighted by arrows in Fig. 2A). In one example, a larger (^~^1.4 μm diameter) circular structure was observed moving over approximately 2 μm, while maintaining its size and the same slightly higher signal compared to the GFP-COL signal from the surrounding ER (Fig. 2C, circle and Video 7), but this structure did not interact with the Golgi. As seen in Figure 2C and 2D, again no notable vesicular transport towards the Golgi can be detected from the timepoint of accumulation around the Golgi to the end of the measurement after 26 mins asc/biotin in this cell (Video 7). Extensive analysis of GFP-COL-RPE also failed to reveal any significant GFP-positive large carriers contributing to the accumulation of GFP-COL in the Golgi. The earliest stages of accumulation of GFP-COL proximal to the Golgi suggest an accumulation of GFP-COL at the ER-Golgi interface (e.g. Fig. 2D and Video 7 t=9’30” – t=11’00”), prior to entering the Golgi.

**Figure 2:**
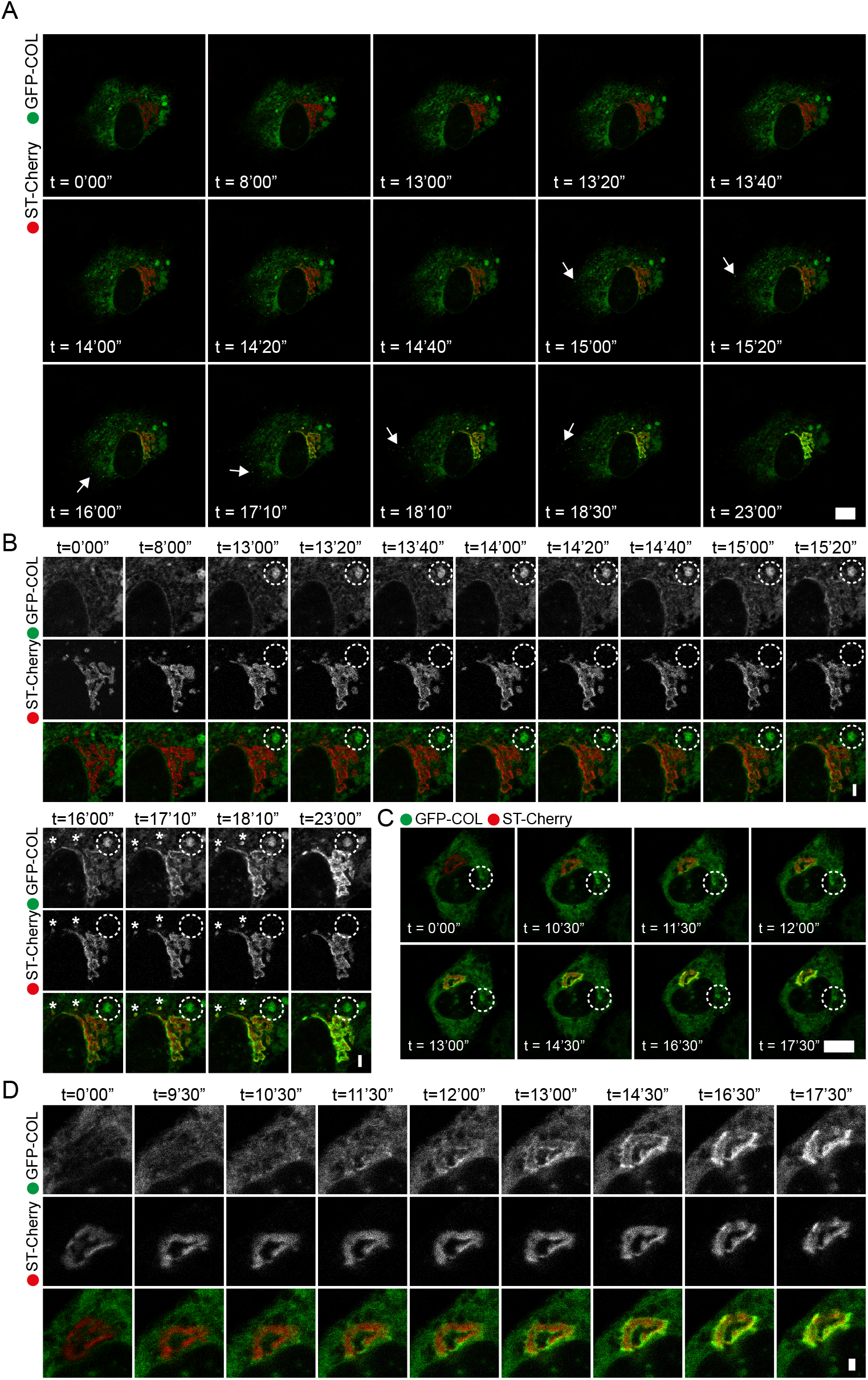
Transport of GFP-COL to the Golgi occurs without the use of large carriers. Image stills from confocal live cell imaging of RPE-1 stably expressing GFP-COL (green; GFP-COL-RPE) co-transfected with the trans-Golgi marker ST-Cherry (red). Timepoints indicate mins after addition of asc/biotin. A: Shows a whole cell imaged at 1 frame every 20 seconds (derived from Video 6). Probable post-Golgi structures are indicated by arrows. Corresponding enlargements of A are shown in B with channels for GFP-COL and ST-Cherry in greyscale, as well as the overlay image below. Large circular GFP-COL-positive structures appear negative for the trans-Golgi (B, circles). Similar smaller structures are positive for GFP and the Golgi marker (asterisks) and follow the previously described phenotype of concentration of GFP-COL at the edge of the Golgi (t = 14 – 18 mins), followed by filling of the Golgi (t = 23 mins). C: Shows image stills of a whole cell taken at 1 frame every 30 seconds (derived from Video 7). Accumulation and filling of the Golgi with GFP-COL occurs within t = 10.5 mins. D: Corresponding enlargements of the Golgi area in C. Channels are displayed separately in greyscale with followed by a merge image. A total of n = 4 sets was acquired. For each set of live imaging experiment 3 cells from the same dish were imaged simultaneously. Scale bars = 10 μm and 1 μm (in enlargements).

### Large GFP-COL structures are positive for ER and Hsp47, not COPII

To further define the nature of the large immobile GFP-positive structures cells were fixed during live cell imaging experiments at time points when an accumulation of GFP-COL in or around the Golgi was observed (Videos 6 and 8). Fig. 3A shows labelling for the endogenous *cis/medial*-Golgi marker giantin and ST-Cherry in these cells; the separation between markers is consistent with the expected localisation of ST-Cherry as a *trans*-Golgi marker. We found that the large GFP-positive structures observed were negative for giantin (Fig. 3Aii, circle), however they did colocalise with the collagen-specific chaperone Hsp47 (Fig. 3Bi, circle). Therefore, these structures are not bona fide Golgi elements. To determine whether these large Hsp47-positive structures were part of the ER or post-ER structures, we used an ER membrane marker, Cyt-ERM-mScarlet-i (ERM). This construct encodes mScarlet-i fused to the ER-targeting sequence of cytochrome P450 (Costantini et al., 2012). Intriguingly, the large GFP-COL structures colocalised with both Hsp47 and ERM (Fig. 3Ci - ii, circles).

**Figure 3:**
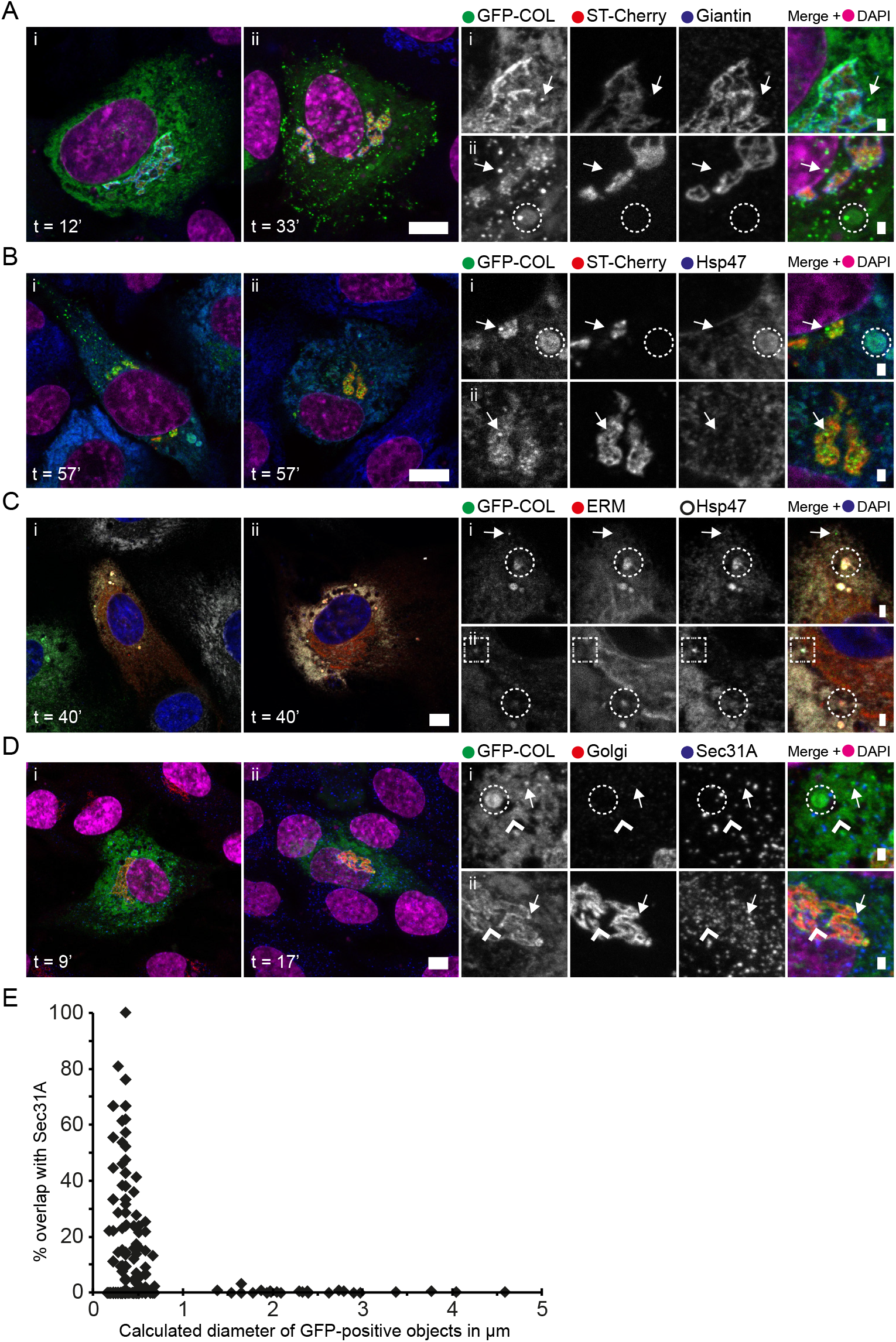
Large GFP-COL structures are positive for ER markers and Hsp47, but not COPII. Confocal imaging of RPE-1 cells stably expressing GFP-COL (green; GFP-COL-RPE) co-transfected with either ST-Cherry (red; A, B, D; 16 - 20 hours post-transfection) or the ER membrane marker ERM-mScarlet-i (red; C; 6 - 8 hours post-transfection). Cells were fixed after the given timepoints (mins after addition of asc/biotin (500 μg.ml-1 and 400 μM, respectively)) and labelled post-fixation with antibodies against further proteins of interest. Large panels show whole cells, while smaller panels show corresponding enlargements of areas of interest with the separate channels in greyscale, followed by the merge image including DAPI (acquired as separate channel and displayed in magenta in A, B and D or blue in C). Scale bars indicate 10 μm and 1 μm (enlargements). n ≥ 10. A: Maximum projection images of z-stacks containing the Golgi apparatus. Duration of asc/biotin corresponds to duration of prior live imaging at approximately one image every 30 seconds until trafficking of GFP-COL to the Golgi was detectable by eye. Cells were labelled with the cis/medial Golgi marker Giantin (blue). Ai (after corresponding Video 8.i) shows an accumulation of GFP-COL at the edge of the Golgi, without visible large GFP-positive structures. Small puncta can be observed in the cell periphery and close to the Golgi (Ai - ii, arrows). Large GFP-COL-positive structures are negative for both the trans- and cis/medial-Golgi markers (Aii, circles). The corresponding video of live imaging prior to fixation of Aii is shown in Video 6 with image stills shown in Fig. 2A - B. B: Duration of asc/biotin corresponds to duration of prior live imaging at approximately one image every 30 seconds until trafficking of GFP-COL to the Golgi was detectable by eye (Video 8.ii). GFP-COL within the ER at 57 mins asc/biotin colocalises with Hsp47 (blue) in the ER (Bi - ii). GFP-COL puncta in the cell periphery and close to the Golgi are negative for Hsp47 and the trans-Golgi marker ST-Cherry (arrows). Cells that show large (>1 μm diameter) GFP-COL-positive structures appear negative for the trans-Golgi, but positive for Hsp47 (Bi, circles). C: Timepoint indicates incubation with 400 μg.ml-1 ascorbate prior to fixation. GFP-COL colocalises with the transiently expressed ERM-mScarlet-i (ERM; red) and Hsp47 (grayscale) in the ER, including large structures (Ci - ii, circles). Small GFP-positive puncta, likely post-Golgi carriers, show no colocalization with ERM or Hsp47 (Ci, arrows), while some small punctate GFP-COL structures are positive for Hsp47, but not ERM (Cii, square). D: Maximum intensity projection images of z-stacks of whole cells. Duration of asc/biotin corresponds to duration of prior live imaging at approximately one image every 30 seconds until trafficking of GFP-COL to the Golgi was detectable by eye (Video 8.iii and 8.iv, respectively). The red channel marked as “Golgi” shows the combined signal from ST-Cherry and antibody-labelling for giantin in the same channel to enable complete visualisation of the Golgi apparatus. Observed large GFP-positive structures do not colocalise with the COPII marker Sec31A (blue; Di, circles). Small GFP-COL puncta (<0.5 μm in diameter) close to the Golgi and in the cell periphery label for Sec31A (Di-ii, arrow heads). Most small punctate GFP-positive structures do not colocalise with Sec31A (Di - ii, arrows). E: Size distribution of GFP-positive objects relative to colocalization with the COPII marker Sec31A from confocal z-stacks with ∆z = 0.29 μm and a suffcient number of slices to represent whole cells (as shown in D). The x-axis shows the calculated object diameter of GFP-positive objects in μm, while the y-axis shows the calculated percentage overlap of GFP-positive objects with Sec31A. Images for analysis were obtained after live cell microscopy as described in Fig. 1 and Fig. 2 and samples were fixed at the time when an accumulation of GFP-COL at the Golgi was visible by eye. In 11 of the 55 analysed cells no punctate or large GFP-COL objects were detected. These cells only show a GFP-COL accumulation around or in the Golgi. The remaining 44 cells show a total number of 1149 detect objects in the GFP-channel.

To investigate whether any of these structures were COPII carriers, cells were co-labelled for the COPII marker Sec31A (Fig. 3D). We analysed the size distribution of GFP-positive structures relative to their overlap with Sec31A (Fig. 3E). Large GFP-COL structures (>1 μm diameter) were found in approximately one fifth of the cells analysed but these did not show any colocalization with Sec31A (Fig. 3Di, circle and Fig.3E); nor did 95% of small GFP-positive structures (<350 nm diameter) (example shown in Fig. 3Di - ii, arrows). Any GFP-positive puncta that did show a higher percentage of overlap with Sec31A (example shown in Fig. 3Di – ii, arrowheads) were consistently smaller than those that were negative for Sec31A (Fig. 3E). Furthermore, the small GFP-positive puncta were negative for both ERM (Fig. 3Ci – ii, arrow and square) and Golgi markers (Fig. 3Ai – ii, arrows). Interestingly, while most small punctate GFP-positive structures were also negative for Hsp47 (Fig. 3Bi - ii, 3Ci, arrows), some were found to indeed colocalise with Hsp47, but not ERM (Fig. 3Cii, square).

In all cases, as shown in Fig. 2D and Fig. 3A, cells show a gradual accumulation of GFP-COL around the rim of the Golgi followed by subsequent filling as described above for Fig. 1D, without the appearance of significant numbers of peripheral carriers of any size directed towards the Golgi. The only structures positive for both GFP-COL and COPII were small and did not stand out in size compared to other COPII puncta (it should be noted that with confocal imaging and the measuring approach used, objects with a diameter ≤160 nm cannot be differentiated in size). We also found that these larger structures do not label for markers of autophagosomes (Cherry-LC3B, ATG16L, and WIPI2, Supplemental Figure S3A-C, circles) or endosomal markers (EEA1, Rab11, or the transferrin receptor, Supplementary Figure S3D-F, circles).

We observed GFP-COL emerging from Str-KDEL-IRES-mScarlet-i-Sec23A (mSc-Sec23A)-positive ER exit sites (Fig. 4A, enlarged in B and video 9). The filling of adjacent Golgi-elements from proximal ERES are apparent in the enlargements shown in Fig. 4B (arrowheads). We further validated the COPII-dependence of this process using a mutant of Sar1 (Sar1-H79G) which prevents GTP-hydrolysis and so inhibits the COPII reaction. Expression of Sar1-H79G blocks GFP-COL in the ER even in the presence of ascorbate and biotin (Fig. 4C).

**Figure 4:**
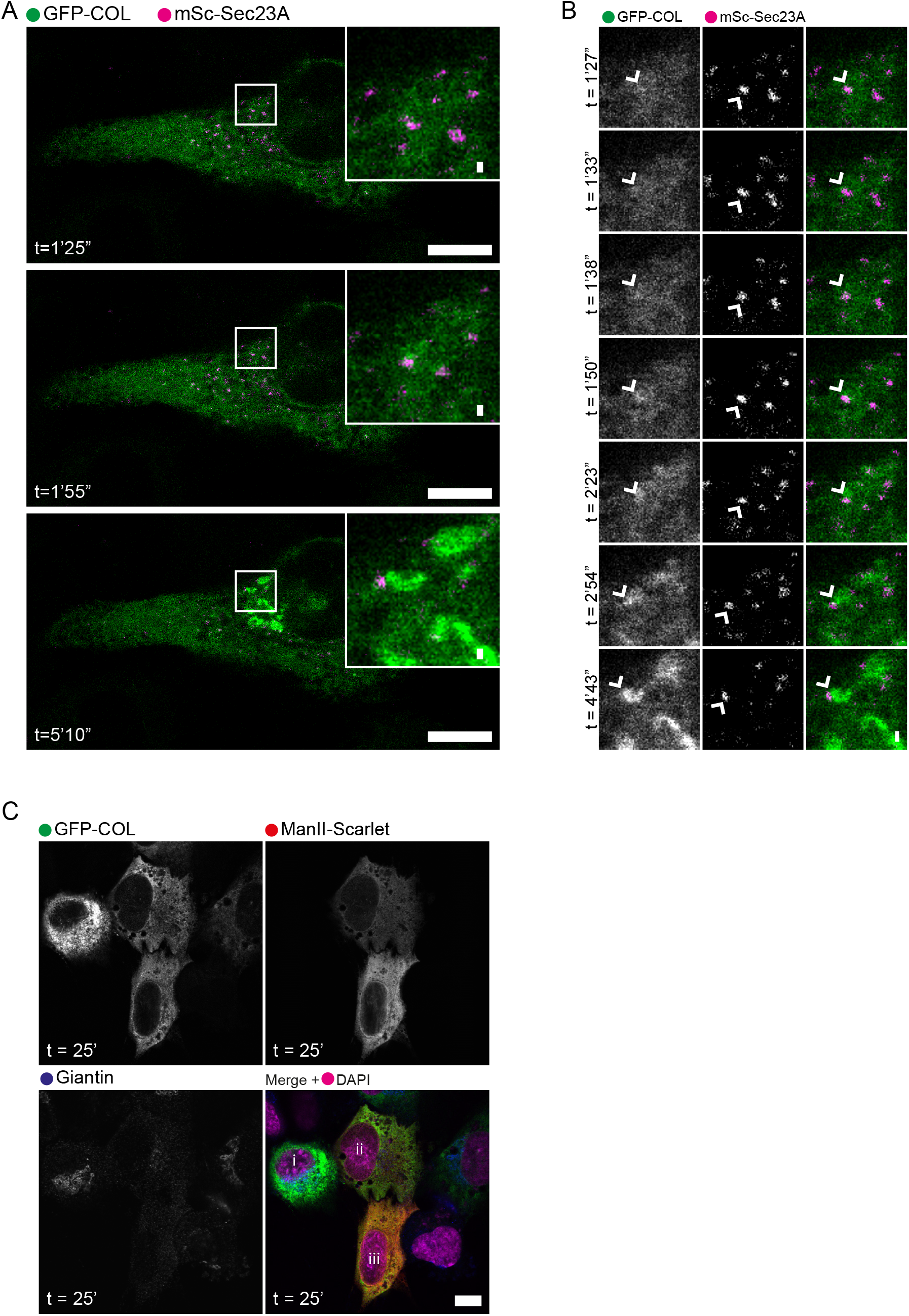
GFP-COL transport is COPII-dependent. A – B: Still images from confocal live cell imaging of RPE-1 stably expressing GFP-COL (green; GFP-COL-RPE) co-transfected with the inner-layer COPII marker mSc-Sec23A (magenta) derived from Video 9. Acquisition at 1 frame every 2.79 seconds. Time points indicate time in presence of asc/biotin (500 μg.ml-1 and 400 μM, respectively) and 20 hours post-transfection. Scale bar = 10 μm (0.25 μm for enlargements) and n = 9. A: Large panels show the entire cell with enlargements displayed in the top right corner. B: Shows the zoomed in area of interest from A (also indicated by the square) with the separate channels in greyscale, followed by the overlay image. Arrow heads indicate a GFP-COL structure that colocalises with a mSc-Sec23A structure and at about 1 min 38 sec before the GFP-COL structure becomes part of the Golgi network. Scale bar = 0.25 μm. C: Example image of a GFP-COL-RPE cells co-transfected with ManII-mSc and the GTP-locked form of Sar1 (Sar1-H79G), which results in a COPII block. Panels show the separate channels GFP-COL (green) and MannII-mSc (red), as well as antibody labelling with a cis-Golgi marker giantin, in greyscale followed by the overlay image including nuclear labelling for DAPI (imaged as a separate channel in pseudo colour magenta). The time point indicates the incubation in presence of asc/biotin prior to fixation with PFA and 7.5 hours after transfection. Scale bar = 10 μm. Cells expressing Sar1H79G show a scattered a disrupted Golgi apparatus, marked by ER-labelling, compared to visible Golgi stacks in cells not affected by the COPII block (i). No transport of GFP-COL can be observed in COPII blocked cells (ii and iii).

To provide better resolution, we used gated stimulated emission depletion microscopy (gSTED) to resolve GFP-COL and COPII-positive structures. Fig. 5A (enlargement in 5B) shows GFP-COL-RPE labelled with both mSc-Sec23A and antibodies against endogenous Sec24C and GFP to maximise detection of the inner layer of the COPII coat and of GFP-COL respectively. We observed GFP-COL-positive structures ranging from <100 nm to >1 μm and quantified the diameter of these objects by fitting the corresponding line graphs (where possible) to a Gaussian function and measuring the full width at half maximum intensity (FWHM). The enlargements, and associated quantification using line scans (Fig. 5C), show clearly that the largest COPII-positive GFP-COL structures (e.g. Fig. 5Ci, 5Cvii and 5Cx) are not uniformly COPII-positive; occasionally they are associated with small COPII puncta but these are not consistent with encapsulation and instead are more consistent with clusters of ERES as we have previously described (McCaughey et al., 2016). Where GFP-COL-positive and COPII-positive structures are seen (e.g. Fig 5Cii, 5Ciii, 5Civ, 5Cv, and 5Cix), they are typically smaller than 350 nm. Furthermore, large circular GFP-positive structures with diameters above 1 μm remain negative for COPII labelling with mSec23A/Sec24C (Fig. 5Cvi and 5Cviii). Therefore, we sought to examine the transport and occurrence of large carriers of endogenous procollagen in fibroblasts secreting large quantities of type I procollagen.

**Figure 5:**
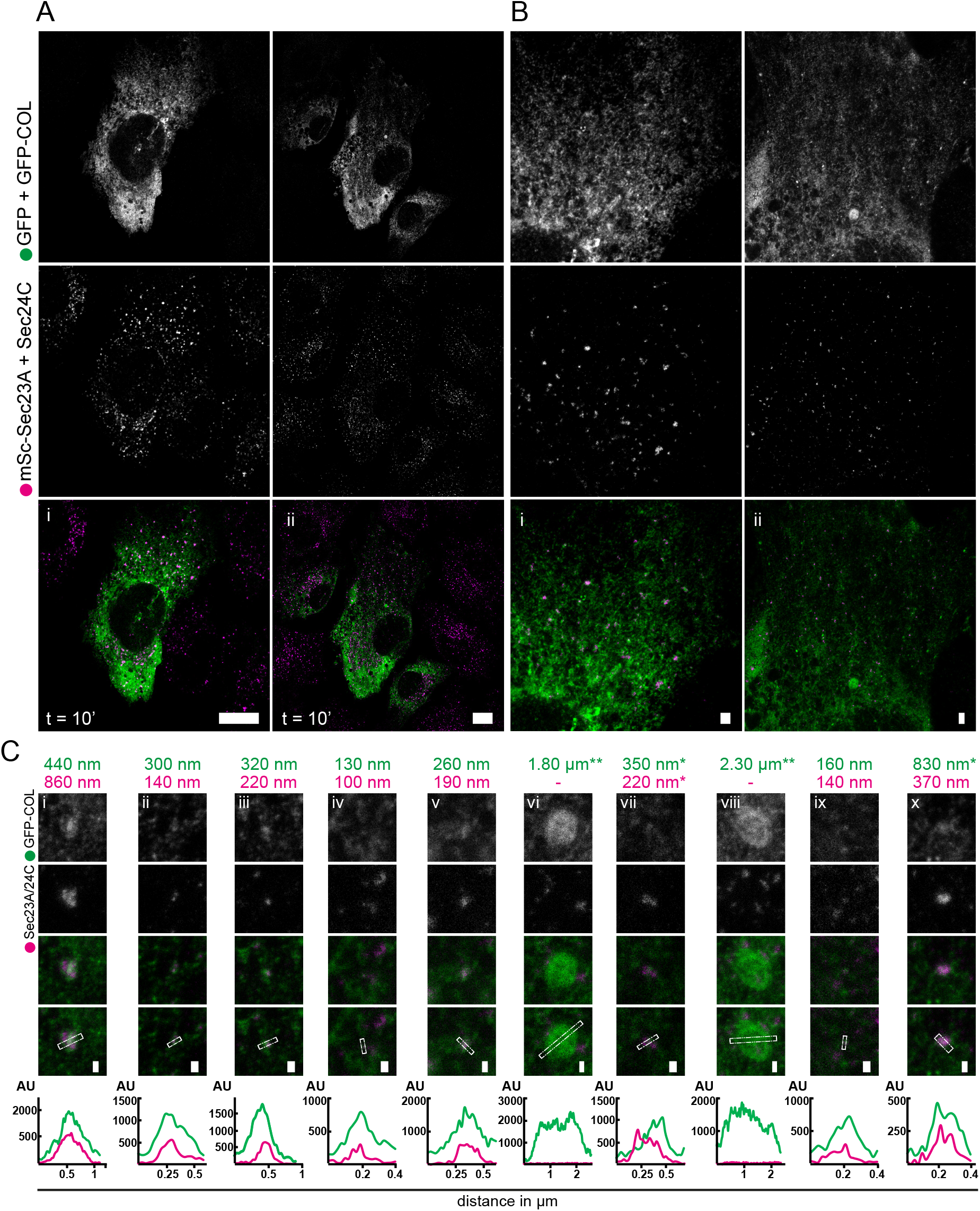
GFP-COL puncta co-labelling for the inner COPII-layer do not exceed 450 nm in diameter. A: Confocal images obtained using the STED microscope of whole RPE-1 cells stably expressing GFP-COL (GFP-COL-RPE; co-labelled with an antibody against GFP in the same channel; green) co-transfected with the inner layer COPII-marker mSc-Sec23A and additionally co-labelled with an antibody against Sec24C in the same channel (magenta). Timepoints indicate incubation in presence of asc/biotin (500 μg.ml-1 and 400 μM, respectively) prior to fixation with PFA. Scale bar = 10 μm and n= 14 B: Corresponding zoomed in areas of the cells in A, obtained using super resolution microscopy (gSTED). Scale bar = 1 μm. C: Enlargements of small GFP-positive puncta co-localising with the COPII marker and large GFP-positive structures extracted from images as displayed in B. Panels show the different channels in greyscale, followed by the overlay image and the overlay image containing the line with a width of 5 (ii – v, vii and ix) or 10 pixels (i, vi, viii and x) drawn through the object of interest to generate the corresponding line graphs, as shown below. Line graphs display the distance in micrometre on the x-axis and the signal intensity as an arbitrary unit (AU) on the y-axis. The FWHM values corresponding to the estimated maximum object diameters in nanometre or micrometre are displayed above the images for the corresponding line graphs fitted with a Gaussian curve (in green for GFP+GFP-COL and in magenta for mSc-Sec23A + Sec24C). When two curves were used to fit the graphs, the sum of the FWHM was used as an estimate*. When Gaussian fitting was not possible, the displayed estimated diameter value was measured via ImageJ**. If no peak in the red channel could be identified, values were left blank (-).

### Primary fibroblasts are devoid of large procollagen structures

Immunofluorescence of endogenous procollagen 1α1 in adult primary skin fibroblasts (NHDF-Ad) showed that it colocalises with the collagen specific chaperone Hsp47 in the ER (Fig. 6Ai and iv). We also observed partial localization to areas of the Golgi apparatus (Fig. 6Aii - iii, arrows), especially at the rim of the Golgi. This appearance is very similar to prior observations made using GFP-COL expressing cells (most evident in Fig.1 Ciii, D (t=40) and Fig.3Ai - ii, 3Bi – ii). However, in primary fibroblasts this observation is independent of ascorbate since colocalization with Golgi markers was observed before, as well as after, 30 mins of incubation with ascorbate. It is possible that sufficient ascorbate was present in the growth medium to facilitate some proline hydroxylation and procollagen export. Importantly, most procollagen-positive punctate structures were negative for the COPII marker Sec31A (Fig. 6Bi - iv, arrows). Similarly, analysis of non-transformed, telomerase immortalised, human fibroblasts (BJ-5ta) showed procollagen accumulated in the Golgi area both in the presence and absence of ascorbate (Fig. 6Ci - iv). Some small procollagen puncta were evident, but again these did not colocalize with Sec31A (Fig. 6Ci - iv, arrows). It was however notable that no large structures labelled for either endogenous procollagen 1α1 or Sec31A were detected in these fixed samples in either cell line. Therefore, our data support a direct route of transport from the juxtanuclear ER to the Golgi without the use of large carriers for both endogenous and overexpressed type I procollagen.

**Figure 6:**
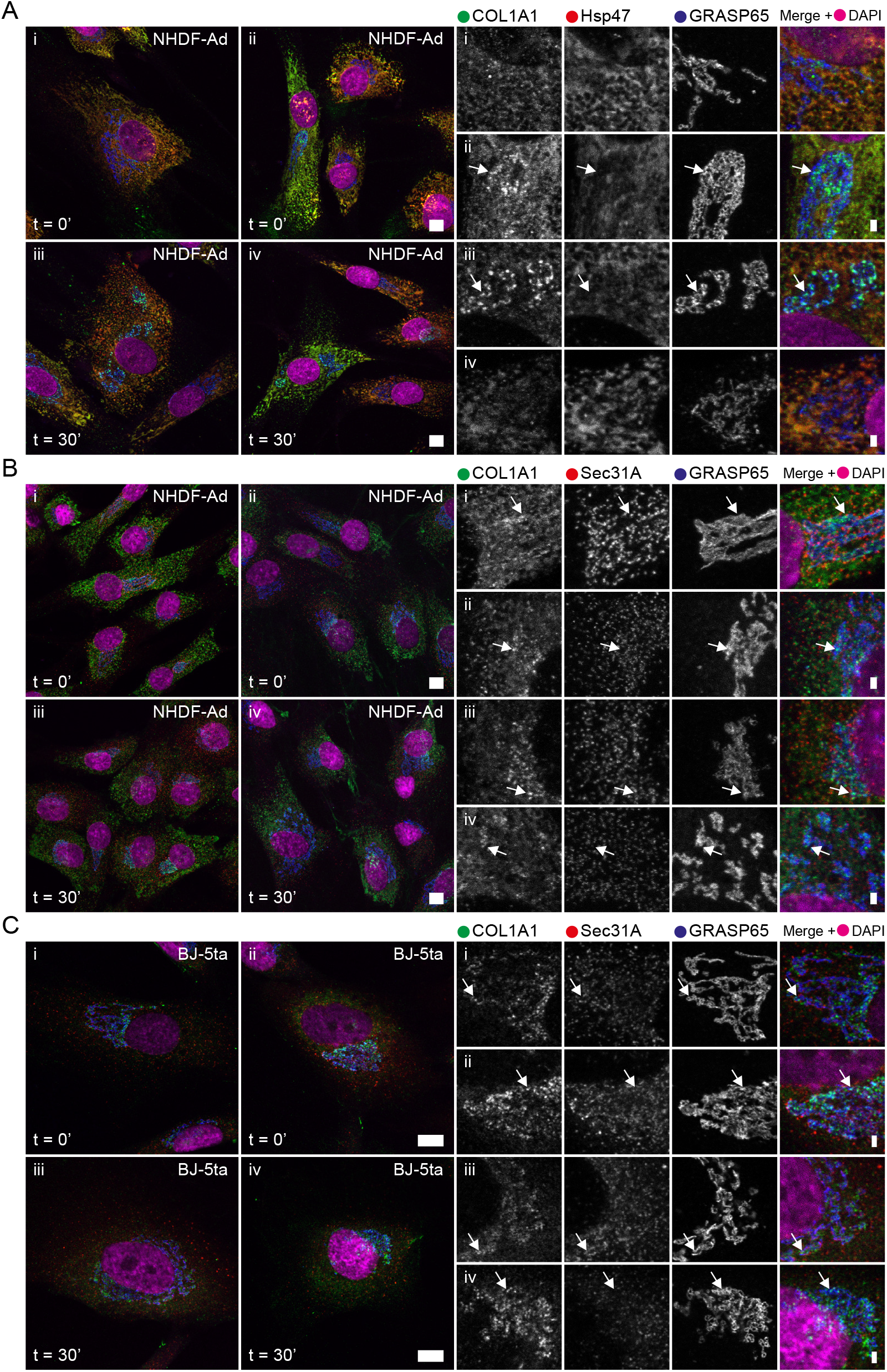
In primary fibroblasts, endogenous procollagen colocalises with Hsp47 in the ER. Confocal images of cells after incubation in the presence of 500 μg.ml-1 ascorbate (A - B) or 50 μg.ml-1 ascorbate (C) for 0 (i - ii) and 30 mins (iii - iv), respectively. A – B: Maximum intensity projection images of z-stacks through primary adult skin fibroblasts NHDF-Ad. C: Shows single z-slice images of non-transformed foreskin fibroblasts (BJ-5ta). Cells were labelled using antibodies against COL1A1 (green) and either Hsp47 (A) or the COPII marker Sec31A (B – C) in red, as well as a cis-Golgi marker GRASP65 (blue). Panels show whole cells, as well as corresponding enlargements of the Golgi area on the right. Enlargements show the separate channels in greyscale as well as the overlay image including nuclear staining in magenta (DAPI, imaged as separate channel). Arrows highlight punctate COL1A1 structures with high signal intensity localising in close proximity to the Golgi. Scale bars = 10 μm for large panels and 1 μm for enlargements. NHDF-Ad show varying expression levels of Hsp47 and COL1A1 (Ai - iv). Localisation of COL1A1 to punctate structures in the Golgi area occurs at both 0 and 30 mins in the presence of ascorbate (Aii, iv and Bi - iv, arrows). BJ-5ta show lower levels of COL1A1 expression compared to NHDF-Ad, but accumulation of COL1A1 in the Golgi area can also be observed at both time points (Ci - iv, arrows). n ≥ 10 in each case.

We also tested a variety of other anti-COL1A1 antibodies for immunofluorescence with NHDF-Ad and IMR-90 cells (Supplementary Fig. S4). None of these cells show any large COPII (Sec24C or Sec24D) and/or COL1A1 structures, even when labelling with the same mouse monoclonal antibody previously shown to visualise large COL1A1 structures by other labs (Gorur et al., 2017).

### GFP-COL transport to the Golgi is not dependent on an intact microtubule network

The absence of long-range carriers translocating from the cell periphery to the Golgi during live cell imaging led us to question the role of peripheral ER exit sites in procollagen transport. Long-range ER-to-Golgi transport in mammalian systems occurs by translocation along microtubules. Microtubules are not essential for trafficking but optimise the efficiency of transfer between organelles. An intact microtubule network is, however, required to maintain the juxtanuclear organization of the Golgi ribbon. Upon microtubule disassembly, such as in the presence of the drug nocodazole, the Golgi scatters into functional mini-stacks dispersed throughout the cytosol adjacent to ERES. Nocodazole can thus be used to disrupt long-range trafficking from peripheral ERES without affecting local ER-Golgi transport at the sites of the mini-stacks.

To determine the requirement for microtubules, and a juxtanuclear Golgi, in the transport of procollagen, cells were treated with nocodazole (NZ) for 60 - 120 mins prior to imaging. Efficacy of nocodazole was validated using tubulin labelling (Supplementary Fig. S5A). Fig. 7 shows images of GFP-COL-RPE cells 60 mins after the addition of NZ followed by addition of asc/biotin/NZ. As expected, the Golgi apparatus redistributed to scattered structures of either circular shape or as short ribbons. In Fig. 7A and 7B GFP-COL is distributed throughout the ER, while two larger GFP-COL structures (diameter of approximately 1.5 – 2 μm) could be seen close to the nucleus with a signal intensity higher than that of GFP-COL in the ER background. In this example, GFP-COL structures are first visible, distributed throughout the cell, 19 mins after addition of asc/biotin/NZ (Video 10). Nearly all structures appeared at the edge of the scattered *trans*-Golgi elements. Gradual accumulation of GFP-COL at the edge of the *trans*-Golgi further increased for about 4 mins. No vesicular-tubular or circular transport carriers could be identified. An even distribution throughout the *trans*-Golgi could be observed between 26 and 35 mins after addition of asc/biotin/NZ (depending on the individual Golgi element) (Fig. 7A – B, Video 10). The overall signal intensity for GFP-COL subsequently decreased over time. Once the *trans*-Golgi had filled, small GFP-COL puncta were seen emerging from Golgi elements and moving towards the cell perimeter. Subsequent emptying of the Golgi, consistent with onward trafficking of GFP-COL, was indistinguishable from traffic in the absence of nocodazole (Fig. 7C, t=44’14”).

**Figure 7:**
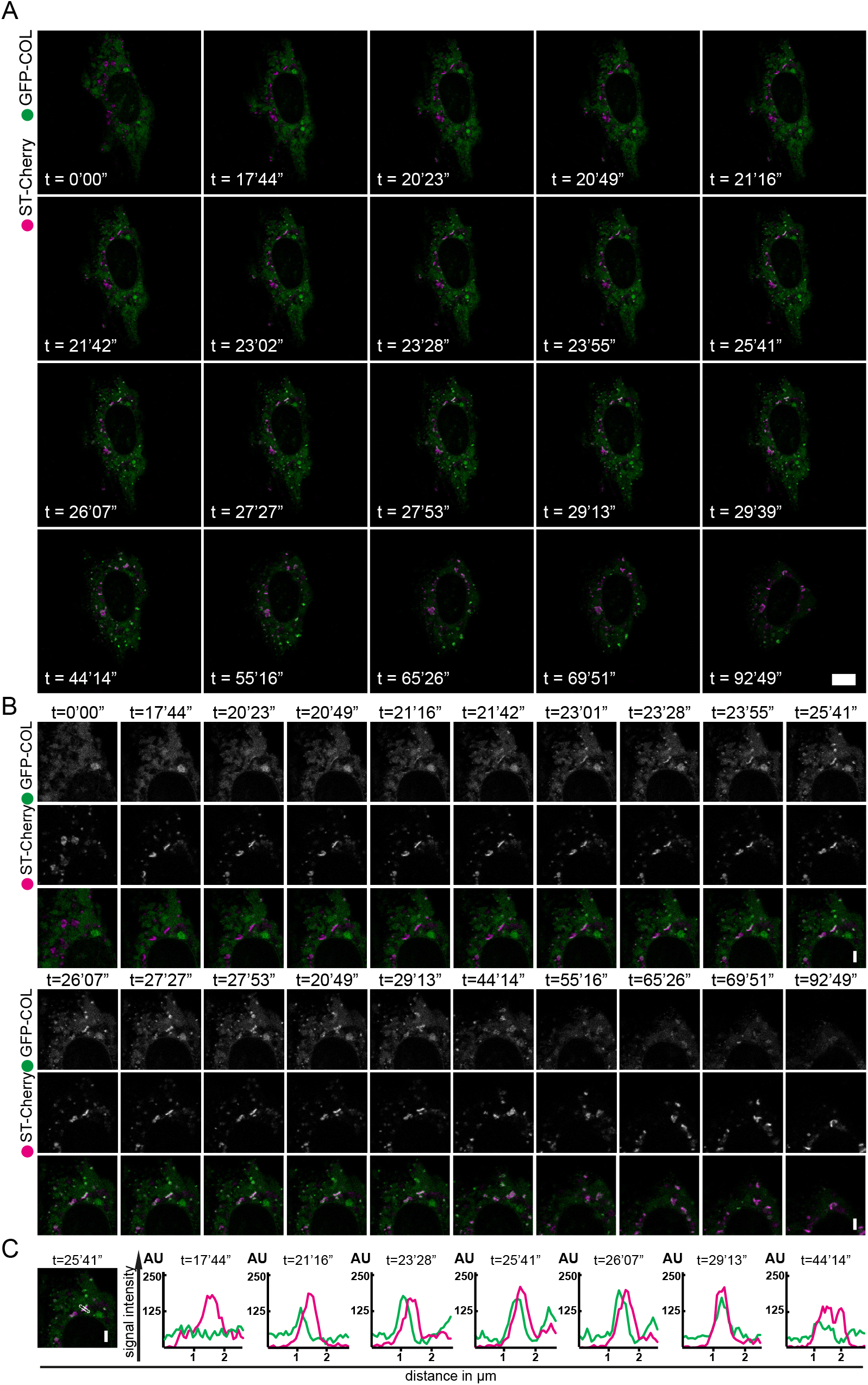
GFP-COL traffcking does not rely on an intact microtubule network. Still images from confocal live cell imaging of RPE-1 stably expressing GFP-COL (green; GFP-COL-RPE) co-transfected with the trans-Golgi marker ST-Cherry (magenta) derived from Video 6. Acquisition at 1 frame every 26 seconds. Cells were incubated in presence of nocodazole for 60 mins prior to imaging. Time points indicate time in presence of asc/biotin (500 μg.ml-1 and 400 μM, respectively). A: Large panels show the entire cell, while enlargements in B highlight a zoomed in area of interest. Small panels show the green channel and magenta channel in greyscale followed by the overlay. Scale bars indicate 10 μm, 1 μm (in enlargements). C: Example image of an enlarged overlay with a 5-pixel wide line drawn through the Golgi followed by corresponding line graphs of selected timepoints showing the signal intensity (y-axis) in arbitrary units for GFP-COL (green) and ST-Cherry (magenta) for the corresponding line. X-axis shows the distance in μm. n = 4.

## Discussion

The currently accepted model for ER export of fibrillar procollagens, such as COL1A1, requires transport in large vesicles (Malhotra and Erlmann, 2015; Miller and Schekman, 2013; Saito and Katada, 2015; Stephens, 2012; Venditti et al., 2014). There are several reports of these structures in the literature from both cell-based and *in vitro* experiments (Gorur et al., 2017; Jin et al., 2012; McGourty et al., 2016; Yuan et al., 2017). Our data, using both stable expression of a newly engineered fluorescent procollagen reporter and endogenous labelling, do not support that model, at least for the long-range translocation of carriers from the peripheral ER to the Golgi. Analysis of fixed cells shows that, where detected, most carriers are <350 nm in size. We also do not detect large structures that are COPII-coated. However, live cell imaging does not support a model where these are a major component of ER-to-Golgi trafficking. In contrast, our work shows that GFP-COL accumulates at sites juxtaposed to the Golgi without the formation of large carriers. Analysis of endogenous procollagen trafficking in fibroblasts confirmed this observation.

The construct we describe here is tagged at the N-terminus before the protease cleavage site. Very recently, similar constructs where the α2 chain of type I procollagen (COL1A2) was tagged at the N-terminus were described (Lu et al., 2018). In that work, the N-terminal propeptide was replaced with a fluorescent protein and the cleavage site removed. This was done intentionally and enabled the visualization of procollagen in the extracellular matrix. In our construct, we chose to retain the cleavage site such that extracellular procollagen itself does not become fluorescent. This enabled us to image intracellular trafficking more effectively. Furthermore, atypical retention of the N-propeptide due to impaired cleavage can lead to pathologies that overlap with those of osteogenesis imperfecta and Ehlers-Danlos Syndrome (Cabral et al., 2005; Malfait et al., 2013). Another recent development has been the engineering of a photoactivatable Dendra2 fluorescent protein into the endogenous locus of COL1A2 (Pickard et al., 2018). Here, Dendra2 was placed after the N-terminal cleavage site such that it is also retain in extracellular collagen fibrils. While this has advantages in terms of endogenous tagging and in analysis of fibril turnover, it does not enhance the ability to control ER export or to selectively analyse intracellular precursors versus extracellular pools.

A major caveat to the use of fluorescent protein-tagged reporters of procollagen is the question of how much of the visualized reporter is indeed trimeric, assembled procollagen. We chose to tag COL1A1 rather than COL1A2 because of its capacity to homotrimerise (Jimenez et al., 1977; Uitto, 1979), potentially lessening the impact of overexpression. We have validated that GFP-COL can and indeed heterotrimerise with COL1A2 in RPE-1 cells using proteomics (Supplementary Figure S5B). However, it should be noted that our own RNAseq data (Stevenson et al., 2017) has shown that COL1A2 is only expressed at very low levels in these cells (shown in Table form in Supplementary Figure S5C). The simplest interpretation is that the COL1A1 homotrimer is therefore the relevant trimer in these cells. We also showed that our stable expression of GFP-COL does not significantly affect expression of COPII proteins including Sec31A, Sec12, TFG, Sec24A, or Sec24C (Supplementary Fig. S5D). In addition, we did not detect major differences in the total amount of COL1A1 expressed by these cells (Supplemental Fig. S5E). We consider it important to note that we, and others, using reporters of this type likely visualize a combination of monomeric and trimeric procollagen exiting the ER. The ascorbate-dependence of ER exit supports the idea that a major pool is indeed trimeric. Regardless, we have not observed large procollagen-containing structures, COPII-coated or otherwise, in any significant number that could mediate the transport of procollagen from the ER-to-Golgi.

Our work supports the possibility of direct connections between the ER and Golgi to facilitate transfer of folded procollagen (Kurokawa et al., 2014; Malhotra and Erlmann, 2015). This idea has parallels with the cisternal maturation model of procollagen Golgi transit, where procollagen does not make use of vesicles while trafficking between Golgi stacks, but rather stays within the cisternae (Bonfanti et al., 1998). Direct connections are difficult to envisage given the well-characterized differences in composition between these organelles. A model that could reconcile this is the local formation of budding structures at ERES in close proximity to Golgi membranes (Fig. 8), as have been generated in *in vitro* budding assays (Gorur et al., 2017; Yuan et al., 2017). Direct connection of the ER to the ERGIC would both prevent compartment mixing of the ER and Golgi and be consistent with data on the role of TANGO1 (Ma and Goldberg, 2016; Nogueira et al., 2014; Raote et al., 2018). One could view such a model as a maturation process where these nascent COPII-coated carriers form the ERGIC itself acquire compartment-specific markers and identity by direct fusion (Fig. 8B). This is also consistent with the nocodazole experiments in which the Golgi is distributed adjacent to peripheral ERES. However, there is no direct evidence of such direct connections or for an equivalent of cisternal maturation from the ERGIC to early Golgi.

**Figure 8:**
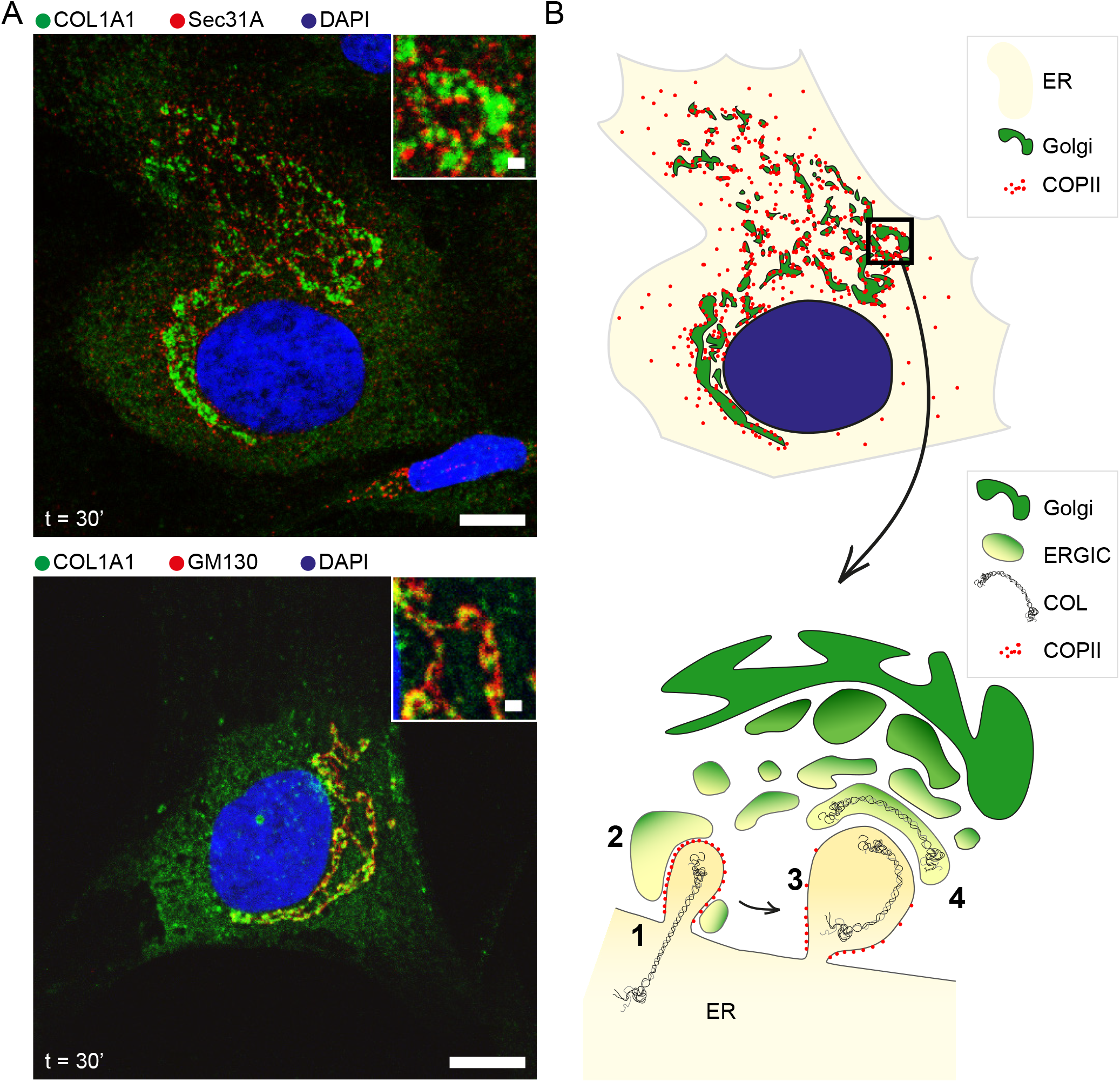
GFP-COL transport to the Golgi via a“short-loop” pathway. A: Immunofluorescence image of a primary skin fibroblast labelled for (i) endogenous COL1A1 (green) and Sec31A (red) and (ii) endogenous COL1A1 (green) and GM130 (red), both after 30 mins incubation in presence with 50 μg.ml-1 ascorbate. B: Schematic of a cell with zoomed in region showing COL1A1 transport from ERES in close proximity to the Golgi. (1) COPII-dependent packaging of procollagen into nascent buds. (2) These buds are expanded by TANGO1-dependent fusion with the ERGIC. (3) Carrier expansion encapsulates procollagen. Questions remain as to the extent of expansion required and the flexibility of the trimer at this stage. (4) Scission of the carrier generates a compartment that almost immediately adopts the identity of the ERGIC and progresses to become the first cisterna of the Golgi. In this model, the ERGIC acts as an intermediate to both expand the nascent ER-derived carrier and to ensure compartmentalization of ER and Golgi.

This model raises questions about how large procollagen molecules could be encapsulated in COPII vesicles. Recent new data have challenged previous assumptions concerning the rigidity of triple helical collagen (Rezaei et al., 2018), showing instead that it can best be described as a semi-flexible polymer. Atomic force microscopy showed that the global curvature of collagen varies greatly with the salt concentration and pH. Importantly, this was seen with many collagen isotypes. It is therefore not inconceivable that the conditions at the point of ER exit are consistent with flexible procollagen that could be packaged into vesicular carriers closer in size to those classically described for COPII.

In many cases, small GFP-COL puncta were seen in our experiments that colocalise with Sec31A. These show a distinct size distribution from large, static, circular structures negative for the COPII marker. Therefore, our data are entirely consistent with COPII-dependent trafficking of procollagen from the ER via conventionally described ER exit sites and we do not dispute an absolute requirement for COPII in this process. Indeed, there is overwhelming support for this from *in vitro* (Gorur et al., 2017; Yuan et al., 2017), cell-based (Stephens and Pepperkok, 2002; Townley et al., 2008), and whole animal experiments (Garbes et al., 2015; Lang et al., 2006; Townley et al., 2008).

Our work also shows that trafficking and secretion of GFP-COL does not require an intact microtubule network. Transport of GFP-COL to Golgi stacks in our experiments appears very similar to that of GFP-tagged tumor necrosis factor (TNF-SBP-EGFP) (Fourriere et al., 2016), supporting the idea that this pathway might not be restricted to procollagen or other large cargo. We also think it unlikely that this short-loop pathway is itself mediated by the formation of large carriers, because large structures of the type described by others (Gorur et al., 2017; Jin et al., 2012; McGourty et al., 2016; Raote et al., 2017) are not evident in our experiments analysing either fixed or live cells. Notably, in those previously published experiments, COPII labelling appears very different to that one might expect from many other studies with few structures evident (Jin et al., 2012). Live imaging of both Sec31A-YFP and procollagen-CFP in KI6 cells that overexpress both procollagen and KLHL12, shows limited movement of an apparent Sec31A-positive and procollagen-positive structure over ~3 minutes (Gorur et al., 2017). This is not entirely consistent with long-range, vectorial ER-to-Golgi transport described previously (Stephens and Pepperkok, 2002) and the highly dynamic nature of the ER itself must be considered. Interestingly, some large FLAG-KLHL12 and Sec31A positive structures in KI6 cells colocalize with Hsp47 (Gorur et al., 2017). These structures would be consistent with our larger GFP-COL structures that are also positive for the collagen chaperone Hsp47.

In our experiments, some larger procollagen-containing structures are visible, but our data shows that these are most likely domains within the ER as they co-label for an ER membrane marker and Hsp47. Being contiguous with the ER, one might expect that these would label with markers of ER exit sites which are normally highly abundant in cells. Notably, we do not observe substantial labelling of these structures with Sec31A. The trafficking of Hsp47 itself is somewhat unclear. While it can bind to both monomeric and trimeric procollagen (Satoh et al., 1996) and has a KDEL retrieval sequence, both consistent with export from the ER, its localization at steady-state is almost entirely restricted to the ER (Duran et al., 2015; Kano et al., 2005; Salo et al., 2016). Current models favour a role for Hsp47 in maintaining the procollagen trimers in a non-aggregated state within the ER (Tasab et al., 2000) and possibly during further transit through the secretory pathway. *In vitro* data show that Hsp47 preferentially binds to the trimeric form of procollagen (Ishikawa et al., 2016; Ito and Nagata, 2017; Koide et al., 2006; Ono et al., 2012; Tasab et al., 2002) and accompanying procollagen after release from the ER (Oecal et al., 2016; Satoh et al., 1996). Interestingly, while most GFP-COL puncta in the cell periphery, as well as those close to the Golgi, were negative for Hsp47, some of these small punctate structures were positive for Hsp47 while also being negative for ER membrane. Therefore, these structures could be bona fide ER-to-Golgi transport carriers.

Mechanistically, a complex machinery, including KLHL12, is clearly involved in COPII-dependent trafficking from the ER. Our data, however, suggest that its role is not to generate large carriers in the cell periphery to direct this traffic. Rather, our findings indicate procollagen trafficking utilises what we term a “short-loop” pathway from the ER to the Golgi. This pathway could of course be used by other cargo. Advances in both light and electron microscopy provide opportunities to address this further. However, we conclude from our work that large carriers do not mediate long-range transport of procollagen from the peripheral ER to the Golgi.

## Materials and methods

Unless stated otherwise, all reagents were purchased from Sigma-Aldrich (Poole, UK).

### DNA constructs

All restriction and modifying enzymes were purchased from New England Biolabs (Hitchin, UK). ER-membrane marker pCytERM_mScarlet-i_N1 was a gift from Dorus Gadella (Addgene plasmid #85068) (Bindels et al., 2017). mCherry-LC3B was a gift from Jon Lane (the GFP-tag from GFP-LC3B ((Betin et al., 2013), pLXG-3 SSFFV GFP-LC3B) was replaced by the Lane lab for a mCherry-tag). To generate the Str-KDEL-IRES-sialyltransferase-mCherry (ST-Cherry; Addgene #110727) construct, Str-KDEL_ST-SBP-mCherry (Addgene #65265), a gift from Franck Perez (Boncompain et al., 2012), was used as a template; the ST-SBP was exchanged for a ST sequence without the SBP-tag via restriction digest with AscI and SbfI. To generate Str-KDEL-IRES-mannosidaseII-mScarlet-i (MannII-mSc; Addgene #117274) and Str-KDEL-IRES-mScarlet-i-Sec23A (mSc-Sec23A; Addgene #117273), Str-KDEL-IRES-mannosidase II-SBP-mCherry (Addgene # 65253), a gift from Franck Perez (Boncompain et al., 2012), and mScarlet-i (Addgene #85044), from Dorus Gadella (Bindels et al., 2017) were used as a template. For MannII-mSc the SBP-mCherry tag was replaced with mScarlet-i. For mSc-Sec23A mannosidaseII-SBP-mCherry were first replaced with mScarlet-i and in a second step Sec23A from pLVX-Puro-mRuby-Sec23A (Addgene #36158; (Hughes and Stephens, 2010)) was amplified using PCR and inserted after Str-KDEL-IRES-mScarlet-i. Sar1-H79G (Aridor et al., 1995).

The GFP-COL construct was designed using the NEB assembly tool (www.neb.com) to introduce a SBP-mGFP-tag between the N-propeptide and the corresponding cleavage site upstream of the triple helical domain of human procollagen 1α1. This was realised by using a two-step NEB HiFi Assembly reaction. First, the inserts procollagen-SBP, a synthetic construct composed of the sequence encoding the signal peptide and N-propeptide of human procollagen 1α1 (synthesised by MWG in pEX-A2), humanised monomeric GFP (mGFP; equivalent to NCBI accession number KP202880.1) and the genetic sequence encoding the triple helical domain and the C-terminus of human COL1A1 (NCBI accession number NM_000088.3) were amplified via PCR (primers listed below). mGFP and COL1A1 were subsequently purified via gel extraction. Prior to assembly, the remaining template vector procollagen-SBP-pEX-A2 was digested using DpnI. The lentiviral vector backbone pLVXPuro (Clontech/Takara Bio Europe, Saint-Germain-en-Laye, France) was linearized via overnight digest with EcoRI and subsequent heat inactivation. For the reaction the compounds to be combined were added in equimolar proportions of 0.05 pmol, except for procollagen-SBP with 0.18 pmol and incubated with the assembly master mix for 3 h at 50 °C, according to the NEB HiFi Assembly protocol. The transformation of NEB 5α competent *E. coli* was performed as described in the protocol. Sequencing of clones after the first assembly reaction showed successful insertion of mGFP and COL1A1 into pLVXPuro and an introduction of a unique PshAI restriction site between pLVXPuro and mGFP, which was used for a second assembly reaction. For this, 0.04 pmol procollagen-SBP, amplified using the primers listed below, and 0.02 pmol of the first assembly product (linearized using PshAI) were used for the assembly reaction. Resulting colonies yielded the desired construct procollagen-SBP-mGFP-COL in pLVXPuro (GFP-COL; Addgene #110726).

Primers (synthesized by Sigma-Aldrich) used for amplification of the inserts: Forward primer for mGFP AGAGCCCATGTGCTGCTGCTGCATGGTGAGCAAGGGCGAG, reverse primer for mGFP GGGGCAGCAGCAGCACTTGTACAGCTCGTCCATGC, forward primer for COL1A1 GCTGTACAAGTGCTGCTGCTGCCCCCAGCTGTCTTATGGC, reverse primer for COL1A1 CGCGGTACCGTCGACTGCAGTTACAGGAAGCAGACAGG and final primers for procollagen-SBP amplification ACTCAGATCTCGACACCGGTCGCCACCATGTTCAGCTTTG (forward) and CCATGGTGGCGACCGGTGTCTTCATGGGCTCTCTCTGG (reverse).

### Cell culture, cell lines and transfection

All cells were cultured at 37 °C and 5 % CO_2_ in a humid environment. IMR-90 (ATCC Cat# CCL-186) were grown in MEM (Sigma-Aldrich) supplemented with L-glutamine and 10% decomplemented FBS (Gibco). DMEM (Gibco) containing 2 % M199 (Gibco) and 10% decomplemented FBS was used for cell culture of BJ-5ta hTERT-immortalized fibroblasts (ATCC Cat# CRL-4001). NHDF-Ad-Der (NHDF-Ad) primary fibroblasts (Lonza Cat# CC-2511) were grown in FBM medium supplemented with FGM-2 (Lonza). For generating the cell line stably expressing the GFP-COL construct, as well as a cell line expressing just GFP as a control, hTERT RPE-1 cell line (ATCC Cat# CRL-4000) were used. These cells were grown in DMEM-F12 Ham (Sigma-Aldrich) supplemented with 10 % decomplemented FBS. Cells were not validated after purchase but were routinely screened annually (and confirmed negative) for mycoplasma contamination.

To make the stable GFP-COL cell line, virus containing the GFP-COL construct was generated using the Lenti-XTM Packaging Single Shots (VSVG) system from Clontech/Takara according to the manufacturer’s instructions (cat# 631275). Growth media was removed from an 80 % confluent 6 cm dish of hTERT-RPE1 and 1 ml harvested virus supernatant supplemented with 8 μg.ml^-1^ polybrene was added to cells. After 1 hour of incubation at 37°C and 5% CO_2_, 5 ml growth media was added. Transfection media was then replaced with fresh growth media after 24 hours. To select for transfected cells, cells were passaged in growth media supplemented with 5 μg.ml^-1^ puromycin dihydrochloride (Santa Cruz Biotechnology, Heidelberg, Germany) 72 hours post transfection. hTERT-RPE1 cells stably expressing GFP (GFP-RPE) were generated previously (Asante et al., 2014). Stable cell lines were maintained in growth media containing 3 μg.ml^-1^ puromycin.

Cells were transfected at 80% confluence according to the manufacturer’s protocol (Invitrogen/Thermo Fisher, Paisley, UK), but using 2.5 μL Lipofectamine 2000 transfection solution (Invitrogen), 1 μg plasmid DNA and 250 μL OPTIMEM per 35 mm well, added drop wise onto the cells with 1 mL of fresh media. Transfection with ST-Cherry, GFP-COL, ManII-mSc or mSc-Sec23A was performed about 16 – 20 hours prior to fixation or live imaging, while transfection with ERM or Sar1-H79G was performed 6-8 hours prior to the procollagen trafficking experiments.

### Immunofluorescence

For immunofluorescence, cells were grown on 13 mm cover slips (0.17 mm thickness #1.5 (Thermo Fisher)) and fixed for 15 min at RT with 4% paraformaldehyde (Thermo Fisher) at RT. For fixation post live imaging, cells were grown in live cell dishes (MatTek Corp, Ashland, MA) and fixed by adding 8% PFA to an equal amount of FluoroBrite DMEM imaging medium (Life Technologies, A18967-01). Cells were permeabilised with 0.1% (v/v) Triton-X100 (Sigma-Aldrich) for 10 mins at RT and blocked with 3% BSA (Sigma-Aldrich) for 30 min. Immunolabelling with primary and secondary antibodies was performed at RT for 1 hour in a humid environment and in the dark. Antibodies as follow were diluted in blocking solution to the final working concentrations or dilutions: 2.5 μg.ml^-1^ for rabbit polyclonal anti-ATG16L (PM040, MBL, Woburn, USA), 0.5 μg.ml^-1^ rabbit polyclonal anti-COL1A1 (NB600-408, Novus Biologicals), 1:250 for rabbit polyclonal anti-N- and C-propeptide of COL1A1 LF39 andLF41 (both gifts from Larry Fisher, NIH (Fisher et al., 1995)), 5 μg.ml^-1^ mouse monoclonal COL1A1-C-terminal antibody 3391 (clone 5D8-G9, MAB3391, Merck, UK), 0.09 μg.ml^-1^ mouse monoclonal anti-N-terminal collagen type I (Sp1.D8, DSHB), 10 μg.ml^-1^ for anti-COL1A1-C-peptide (PIP) #42024 and #42043 (QED Biosciences, San Diego, California, USA), 1.25 μg.ml^-1^ for mouse-monoclonal anti-EEA1 (Clone 14/EEA1 (RUO) #610456, BD Transduction Laboratories), 0.5 μg.ml^-1^ mouse monoclonal anti-GFP (MMS-118P, Covance (now BioLegend), San Diego, USA), 1:2000 for rabbit polyclonal anti-giantin (Poly19243, Biolegend, London, UK), 0.25 μg.ml^-1^ mouse monoclonal anti-GM130 (610823, BD Biosciences, Wokingham, UK), 1:1500 sheep polyclonal anti-GRASP65 (gift from Jon Lane), 0.75 μg.ml^-1^ mouse monoclonal anti-Hsp47 (M16.10A1, ENZO, Exeter, UK), 2 μg.ml^-1^ for rabbit polyclonal anti-Rab11 (R5903, Sigma), 1:250 for rabbit polyclonal anti-Sec24C (Townley et al., 2008) and 1:100 for Sec24D (Palmer et al., anti-transferrin receptor (TfR; clone H68.4, 13-6800, Thermo Fisher Scientific) and 2.5 μg.ml^-1^ for mouse-monoclonal anti-WIPI2 (clone 2A2, MCA5780GA, Bio-Rad,UK).

Samples were rinsed three times with PBS for 5 mins after incubation with primary and secondary antibodies, respectively. As secondary antibodies 2.5 μg.ml^-1^ donkey anti-rabbit Alexa-Fluor-568-conjugated, donkey anti-mouse Alexa-Fluor-647-conjugated or donkey anti-sheep Alexa-Fluor-488-conjugated antibodies were used. For labelling of endogenous procollagen donkey anti-rabbit Alexa-Fluor-488-conjugate was used.

Samples were washed with deionised water and mounted using ProLong Diamond Antifade with DAPI (Invitrogen) for confocal imaging or MOWIOL 4-88 (Calbiochem, Merck-Millipore, UK) mounting media for widefield imaging. Samples fixed after live imaging were incubated with DAPI (Invitrogen) for 3 mins at RT prior to repeated washing and storage in PBS at 4 °C in the dark until further imaging.

To show the effective disruption of microtubules after incubation with 5 μM nocodazole (NZ) for 60 and 120 mins, GFP-COL-RPE were fixed with −20°C methanol for 4 mins, prior to blocking and immunofluorescent labelling with mouse monoclonal anti-α-tubulin antibody (1:1000, clone B-5-1-2, T5168) and subsequent steps as described above.

### Procollagen trafficking experiments and image acquisition

GFP-COL-RPE1 were incubated in medium containing 50 μg.ml^-1^ L-ascorbic acid-2-phosphate (Sigma-Aldrich; termed ascorbate throughout) for 24 hours - 48 hours prior to live image acquisition or fixation, to “flush out” overexpressed GFP-COL before a controlled accumulation in the ER for 24 h. Finally, to synchronise procollagen trafficking in GFP-COL-RPE, medium was supplemented to contain 500 μg.ml^-1^ ascorbate during live imaging or prior to fixation. This higher concentration was found to more effectively synchronise export. Experiments performed with cells co-transfected with an ER-hook (ST-Cherry, MannII-mSc or mSc-Sec23A, to utilise the RUSH system) were conducted 16 - 20 hours post-transfection and the medium contained 500 μg.ml^-1^ ascorbate, as well as 400 μM biotin (Sigma-Aldrich), to enhance synchronisation and trigger GFP-COL release in a time-dependent manner. For analysis of the dependence on the microtubule network, cells were treated as mentioned above, but incubated in presence of 5 μM NZ for 60 – 120 mins prior to the initiation of the trafficking experiment via addition of asc/biotin/NZ to the cells.

Images of cells transiently expressing GFP-COL were obtained through widefield microscopy using an Olympus IX-71 inverted microscope (Olympus, Southend, UK) combined with Exfo (Chandlers Ford, UK) Excite xenon lamp illumination, single pass excitation and emission filters combined with a multipass dichroic (Semrock, Rochester, NY) and images captured on an Orca-ER CCD (Hamamatsu, Welwyn Garden City, UK). The system was controlled using Volocity (v. 5.4.1 Perkin-Elmer, Seer Green, UK). Chromatic shifts in images were registration corrected using TetraSpek fluorescent beads (Invitrogen/Thermo Fisher).

All other images were obtained using confocal microscopy using Leica SP5II for fixed or Leica SP8 for live samples (Leica Microsystems, Milton Keynes, UK). Z stacks of fixed samples labelled with Sec31A were acquired with ∆z = 0.29 μm. For live cell imaging FluoroBrite DMEM (Thermo Fisher) was used as imaging medium. Live cell imaging was performed using a Leica SP8 confocal laser scanning microscope with 63x HC OL APO CS2 1.42 numerical aperture (NA) glycerol lens and an environmental chamber at 37°C with CO_2_ enrichment and Leica LAS X software. Fluorophores were excited using ≤ 2% energy of the 65 mW Argon laser at 488 nm for the green and a 20mW solid state yellow laser at 561nm for the red channel, respectively. Time courses were acquired using the sequential scanning mode between lines, imaging speed set to 700 Hz, two times zoom and detection of the green and the red channel using ‘hybrid’ GaAsP detectors and corresponding notch filters. Each frame was acquired with a three times line average. One to three cells per sample were chosen that showed low to moderate expression of both GFP-COL and ST-Cherry (estimated by eye) and imaged using multi-position acquisition with ‘Adaptive Focus Control’ active for each cycle and each position to correct axial drift between frames. Minimisation of time between frames was undertaken to allow the highest temporal resolution possible with the given positions per sample resulting in second was accomplished by using a single position imaged at two times line averaging, imaging speed set to 1000 Hz and the highest zoom factor possible to capture a whole cell.

All images shown were not altered from the raw data, unless mentioned. Movies and movie stills were enhanced in brightness and contrast for both channels, by using ImageJ autocorrection (except Fig. 1D). All movies including imaging of ST-Cherry or MannII-mSc were registration corrected using ImageJ’s in-built 3D-correction for drifting, applied using the Golgi channel. All movies were smoothed using ImageJ’s in-built *smooth* processing function.

### Super-resolution microscopy

GFP-COL-RPE were co-transfected with mSc-Sec23A and post-24h in presence of 50 μg.ml^-^1 ascorbate followed by 24 hours ascorbate starvation and addition of 500 μg.ml^-1^ asc and 400 μM biotin 10 mins before fixation with PFA as described previously. Samples were labelled with antibodies against GFP (mouse), as well as anti-Sec24C (rabbit). Donkey anti-rabbit Alexa-568-conjugated antibody and donkey anti-mouse Alexa-488-conjugated secondary antibodies were used, and samples mounted with Prolong Diamond without DAPI (Thermo Fisher Scientific).

Super-resolution images were obtained using a gated stimulated emission depletion microscope (gSTED) Leica SP8 X (Leica Microsystems, Milton Keynes, UK). Samples were imaged at 400 Hz scan speed at unidirectional scanning and detected using gated ‘hybrid’ SMD GaAsP detectors and a 100x HC PL APO CS2 oil lens with numerical aperture 1.4 (serial no 506378). Images were acquired at three to five times zoom with a pixel size of 23 - 24 nm. The fluorophores were excited using a white light laser at 488 and 568 nm. Channels were acquired using a sequential setting starting with the red channel (excitation at 568 nm utilizing a 660 notch filter and emission detection set to 578 – 634 nm with 0.3 to 6 nanoseconds gating), followed by the green channel (excitation at 488 nm and emission detection of 500 – 545 nm with gating set to 1.5 – 7.6 nanoseconds) utilizing a 488 notch filter, both using a pinhole od 1 airy unit. STED laser intensities were at 1.424 for the 592nm depletion laser (for the green) and 1.323 W for the 660 nm depletion laser (for the red channel). For both channels frame averaging was set to 3 and line accumulation to 2 using a Leica LAS X 3.4.0.18371 software.

### Data analysis

Localisation analysis of images of GFP-COL-RPE co-transfected with ST-Cherry and antibody labelled against Sec31A (post-fixation after live-imaging until an accumulation of GFP-COL in the Golgi was visible) was performed automatically. Spatial overlap between Sec31A and GFP-COL was measured using a custom plugin for ImageJ/Fiji (Schindelin et al., 2012; Schneider et al., 2012). First, the punctate Sec31A structures were detected using the Fiji plugin TrackMate (Tinevez et al., 2017), fit with a 2D ellipsoidal Gaussian distribution and false identifications isolated and removed according to a low-pass ellipticity filter. GFP-COL, visible in a separate imaging channel, exhibited a mixed distribution of punctate and broader objects, which were identified separately then combined. Images were processed prior to object identification to enhance the respective structures being detected. For punctate GFP-COL structures, images were convolved with a 3D Gaussian kernel to remove noise then processed with a rolling ball filter (Sternberg, 1983) to subtract non-punctate structures. Punctate GFP-COL structures were subsequently identified using the same approach as for Sec31A, but with an additional low-pass sigma (spot width) filter. For broader GFP-COL structures the raw image was also convolved with a 3D Gaussian kernel and rolling-ball filter, albeit with a larger radius. The images were then processed with a 2D median filter and thresholded using the maximum entropy approach (Kapur et al., 1985). Objects were identified as contiguous regions in the binarised image and filtered using a high-pass size filter. At this point punctate and broad GFP-COL structures were combined, with instances of spatial overlap resolved in favour of broad objects, unless the number of punctate objects per broad object exceeded a user-defined threshold of five. GFP-COL also accumulates around the Golgi. To remove these approach to the broader GFP-COL structures, except using the isodata thresholding approach (Ridler and Calvard, 1978). Any GFP-COL structures within 0.5 μm of a Golgi are removed. Finally, GFP-COL structures are filtered based on their mean intensity in the GFP-COL channel. Pixel-based overlap of Sec31A and GFP-COL is calculated along with the area of each object projected into the XY plane.

The code for data analysis is included as a .zip file in supplemental material. The plugin and the source code are also publicly-accessible on a new GitHub repository (https://github.com/SJCross/ModularImageAnalysis). This is linked to a service called Zenodo which provides a permanent DOI reference to that specific version (https://doi.org/10.5281/zenodo.1252337). The files are as follows: "Modular_Image_Analysis-v0.3.2.jar" is the plugin, which now includes all the third party libraries that were previous stored in the /jars folder; "ModularImageAnalysis-v0.3.2.zip" is the source code for the plugin itself; "Analysis.mia" is the .mia workflow file used for the final analysis; "Installation and usage.txt" describes how to install the plugin and run the Analysis.mia analysis file; "LICENSE.txt" is the license for the plugin itself; "dependencies.html" is an HTML formatted page listing all the dependencies used by the plugin and their associated licenses. The analysis was performed with n=4 independent data sets that contained a total of 55 cells.

For estimating diameter sizes of larger GFP-positive objects and small puncta, as well as the closely or co-localising objects labelling for the inner COPII layer (via Sec24C antibody labelling and mSc-Sec23A in the same channel), whole cells were captured using the confocal mode of the STED, prior to zooming in on an area of interest (showing colocalisation and or bigger GFP-positive structures). Using ImageJ’s line tool a line with a width of 5 pixels for smaller objects, and 10 pixels for larger objects, was drawn through objects of interest and line graphs were generated for both channels. Resulting line graphs were subsequently fitted to 1 - 2 Gaussian curves when possible using MatLab R2016a (MathWorks) and the full width at half maximum (FWHM) was obtained and calculated to show the FWHM in nanometres. Where multiple peaks existed, the summed FWHM of both fitted gaussian curves was displayed in the figure instead. For large objects where gaussian fitting was not possible, the diameters measured manually via ImageJ were selected as estimated diameter. The estimated diameters represent the maximum object diameter of the measured objects. A total of 14 cells were investigated and 20 objects of interest measured.

### Analysis of protein and RNA abundances

For the GFP-trap Chromotek GFP-Trap_A (GFP-Trap^®^ coupled to agarose beads particle size ^~^ 90 μm; Code gta-20) were used. GFP-COL-RPE and GFP-RPE (used as control) were seeded near confluent on 15cm dishes and incubated in presence of ascorbate (50μg.mL^-1^) for 24 hours, followed by ascorbate starvation for 24 hours, followed by 15 mins incubation in presence or absence of 500μg.mL^-1^ ascorbate prior to cell lysis. All following steps were performed at 4°C. The lysis buffer contained 10mM Tris-HCl pH 7.4, 50mM NaCl, 0.5mM EDTA, 0.5% (v/v) IGEPAL and protease inhibitor cocktail (Calbiochem). For cell lysis, cells were rinsed with ice cold PBS and incubated in 0.5 mL lysis buffer for 15 mins. Lysates were subsequently collected and incubated for 30 mins while gently mixing. Beads were equilibrated with lysis buffer (without protease inhibitor cocktail and IGEPAL; referred to as dilution buffer) using 20 μl of bead slurry per sample. Lysates collected after centrifugation at 13500 rpm for 10 mins were incubated with the GFP-trap beads for 2 hours. Samples were subsequently centrifuged at 2700xg for 2 mins to collect the beads with bound sample. After washing twice with dilution buffer containing protease inhibitor cocktail, samples were boiled after addition of 43 μl 2x LDS buffer containing a reducing agent (Invitrogen) at 95°C for 10 mins and beads separated from the denatured protein samples by centrifugation as described above. The entire sample was loaded onto a 3-8% Tris-Acetate precast gel (NuPAGE ^®^) and run for 135 mins at 100V in Tris-Acetate running buffer supplemented with antioxidant (All Invitrogen).

Gel slices were subsequently cut and digested for proteomics analysis via mass spectrometry.

Each gel lane was cut into 6 slices and each slice subjected to reduction (10 mM DTT, 56°C for 30 mins), alkylation (55 mM iodoacetamide, room temperature for 20 mins) and in-gel tryptic digestion (1.25ug trypsin per gel slice, 37°C overnight). The resulting peptides were fractionated using an Ultimate 3000 nano-LC system in line with an LTQ-Orbitrap Velos mass spectrometer (Thermo Scientific). In brief, peptides in 1% (v/v) formic acid were injected onto an Acclaim PepMap C18 nano-trap column (Thermo Scientific). After washing with 0.5% (vol/vol) acetonitrile 0.1% (v/v) formic acid peptides were resolved on a 250 mm × 75 μm Acclaim PepMap C18 reverse phase analytical column (Thermo Scientific) over a 150 min organic gradient, using 7 gradient segments (1-6% solvent B over 1mins, 6-15% B over 58mins, 15-32%B over 58 mins, 32-40%B over 5mins, 40-90%B over 1mins, held at 90%B for 6 mins and then reduced to 1%B over 1 min) with a flow rate of 300 nl.mins^−1^. Solvent A was 0.1% formic acid and Solvent B was aqueous 80% acetonitrile in 0.1% formic acid. Peptides were ionized by nano-electrospray ionization at 2.1 kV using a stainless-steel emitter with an internal diameter of 30 μm (Thermo Scientific) and a capillary temperature of 250°C. Tandem mass spectra were acquired using an LTQ-Orbitrap Velos mass spectrometer controlled by Xcalibur 2.1 software (Thermo Scientific) and operated in data-dependent acquisition mode. The Orbitrap was set to analyse the survey scans at 60,000 resolution (at m/z 400) in the mass range m/z 300 to 2000 and the top twenty multiply charged ions in each duty cycle selected for MS/MS in the LTQ linear ion trap. Charge state filtering, where unassigned precursor ions were not selected for fragmentation, and dynamic exclusion (repeat count, 1; repeat duration, 30 seconds; exclusion list size, 500) were used. Fragmentation conditions in the LTQ were as follows: normalized collision energy, 40%; activation q, 0.25; activation time 10 ms; and minimum ion selection intensity, 500 counts.

The raw data files were processed and quantified using Proteome Discoverer software v1.4 (Thermo Scientific) and searched against the UniProt Human database (downloaded 14/09/17; 140000 sequences) plus the GFP sequence using the SEQUEST algorithm. Peptide precursor mass tolerance was set at 10ppm, and MS/MS tolerance was set at 0.8 Da. Search criteria included carbamidomethylation of cysteine (+57.0214) as a fixed modification and oxidation of methionine and proline (+15.9949) as variable modifications. Searches were performed with full tryptic digestion and a maximum of 3 missed cleavage sites were allowed. The reverse database search option was enabled, and all peptide data was filtered to satisfy false discovery rate (FDR) of 1%.

RNA-seq data from WT RPE-1 cells was derived from our own previously published data (Stevenson et al., 2017). Raw RNA-seq data are available in the ArrayExpress database under accession number E-MTAB-5618.

For semi-quantitative analysis of protein levels of Sec24A, Sec24C and COL1a1 levels in WT RPE-1, GFP-RPE and GFP-COL-RPE cells, cells were seeded on 10cm dishes and grown for 4 days. Cells were incubated in 2 mL serum free culture medium supplemented with (Sec24C and D lysates, as well as COL1A1) or without 50 μg.mL^-1^ ascorbic acid for 24 hours. The medium was collected, and the cells lysed for 15 minutes in buffer containing 50 mM Tris-HCl, 150 mM NaCl, 1% (v/v) Triton-X-100, 1% (v/v) protease inhibitor cocktail (Calbiochem) at pH 7.4 on ice. Protein fractions of medium and lysate were centrifuged at 13500 rpm at 4 °C for 10 min. The cell pellet was discarded. The supernatant was denatured and run under reducing conditions on a 3 - 8% Tris-Acetate precast gel (NuPAGE^®^) for 135 mins at 100V in Tris-Acetate running buffer supplemented with antioxidant. Transfer of protein bands onto a nitrocellulose membrane was performed at 15 V overnight. The membrane was blocked using 5% (w/v) milk powder in TBST for 30 minutes at RT and incubated with antibodies against COL1A1 (Novus Biologicals; 0.5 μg.mL^-1^), Sec24A (Satchwell et al., 2013) and Sec24C (1:100;) and 0.5 μg.mL^-1^ mouse monoclonal anti-GAPDH (AM4300, Thermo Fisher Scientific) as loading control for 1.5 hours at RT. After repeated rinsing with TBST, the membrane was incubated for 1.5 hours at RT with HRP-conjugated antibodies diluted in the blocking solution (1:5000) against mouse and rabbit, respectively (Jackson ImmunoResearch). The wash step was repeated, and detection was performed using Promega WB-ECL reaction reagents and autoradiography films with overnight exposure and subsequent development. Cells from the Sec31A, Sec12 and TFG blots were incubated in normal culture medium prior to cell lysis and scraping. The immunoblot for Sec31A was done as described above, while Western Blots showing intracellular levels of Sec12(a gift from Balch lab, (Weissman et al., 2001)) and TFG (0.05 μg.mL^-1^, NBP2-24485, Novus Biologicals) were run on a 4 - 12 % Bis-Tris (NuPAGE^®^) precast gel at 200 V in MOPS running buffer for 50 mins instead.

## Author contributions

Conceptualization: DJS. Methodology: DJS, NLS, SC, JM. Software: SC. Validation: JM, NLS. Formal analysis: JM, NLS, SC. Investigation: JM, NLS. Resources: DJS, SC. Data curation: JM, NLS, SC, DJS. Writing – original draft: JM, SC, DJS. Writing – Reviewing and Editing: JM, NLS, DJS. Visualization: JM, DJS. Supervision: DJS. Project Administration: DJS. Funding Acquisition: DJS.

## Acknowledgments

This work was funded by the Medical Research Council UK (grant no. MR/P000177/1) and the University of Bristol postgraduate research scholarship. We thank the Wolfson Foundation and University of Bristol for funding of the Wolfson Bioimaging Facility. This work also benefited from additional equipment funded by the BBSRC (through BrisSynBio, a BBSRC/EPSRC-funded Synthetic Biology Research Centre, grant L01386X) and an ALERT13 capital grant (BB/L014181/1). We thank other members of the lab for helpful discussions throughout the project. Thanks to Kate Heesom for her help with the proteomics experiments. We thank Franck Perez, Bill Balch, Larry Fisher, and Dorus Gadella for sharing reagents with us and Ash Evans for creating the ST-Cherry construct. We are very grateful to Jon Lane and other members of our lab for reagents and helpful discussions.

## Supplemental Information

Supplementary information including 5 Figures and 10 videos is included.

---

**Supplemental Figure S1:**
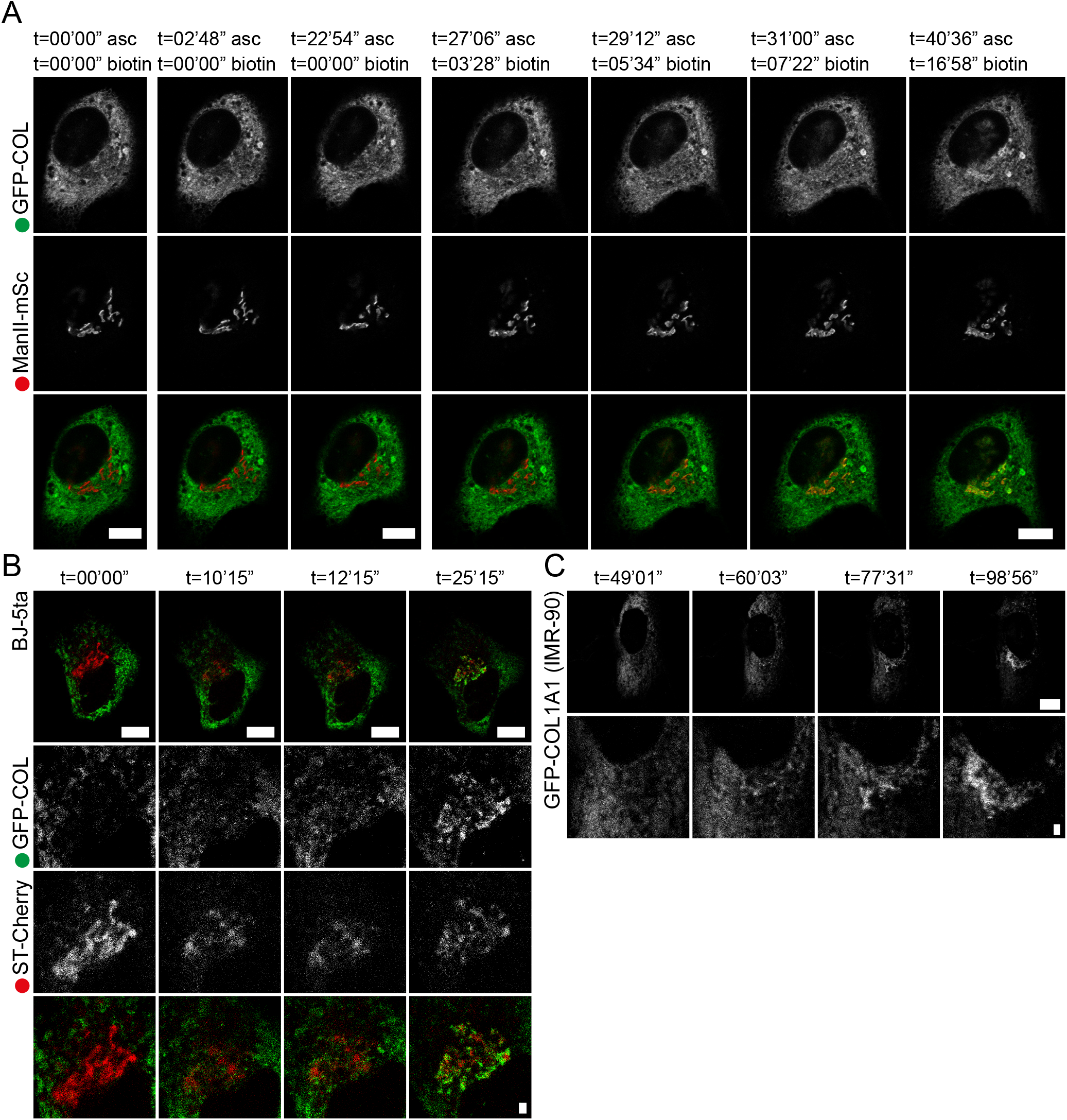
GFP-COL transport is ascorbate and biotin controllable. Image stills from confocal live cell imaging. Scale bars = 10 μm. A: RPE-1 stably expressing GFP-COL (green; GFP-COL-RPE) 17 hours post-co-transfection with the cis-Golgi marker MannII-mSc (red; utilising the RUSH system). Time points indicate total time after addition of 500 μg.ml-1 ascorbate (asc) and 400 μM biotin. Shows a whole cell imaged at 1 frame every 18 seconds (derived from Video 2) with channels for GFP-COL and MannII-mSc in greyscale, followed by the overlay image below. No accumulation of GFP-COL in the Golgi region is visible after 27 mins in presence of ascorbate but appears after about 5 – 7 mins after addition of biotin. A total of n = 2 was acquired. B: BJ-5ta transiently expressing GFP-COL (green) and the trans-Golgi marker ST-Cherry (red; utilising the RUSH system) 17 hours post-co-transfection. Time points indicate time after addition of asc/biotin. The cell was imaged at 1 frame every 15 seconds (derived from Video 3). Panels show the whole cell as overlay with corresponding enlargements of the Golgi area in greyscale for the separate channels GFP-COL and ST-Cherry, followed by the overlay image. An accumulation and filling of the Golgi with GFP-COL becomes visible at about 10 mins after addition of biotin. Scale bars = 10 μm and 1 μm for enlargements. C: IMR-90 cells transiently expressing GFP-COL were imaged 20 hours post-transfection at 2.5 seconds per frame (derived from Video 4). Time points indicate time in presence of ascorbate. An accumulation in the Golgi area is visible at about 60 mins after addition of 500 μg.ml-1 ascorbate. A total of n = 1 was acquired.

**Supplemental Figure S2:**
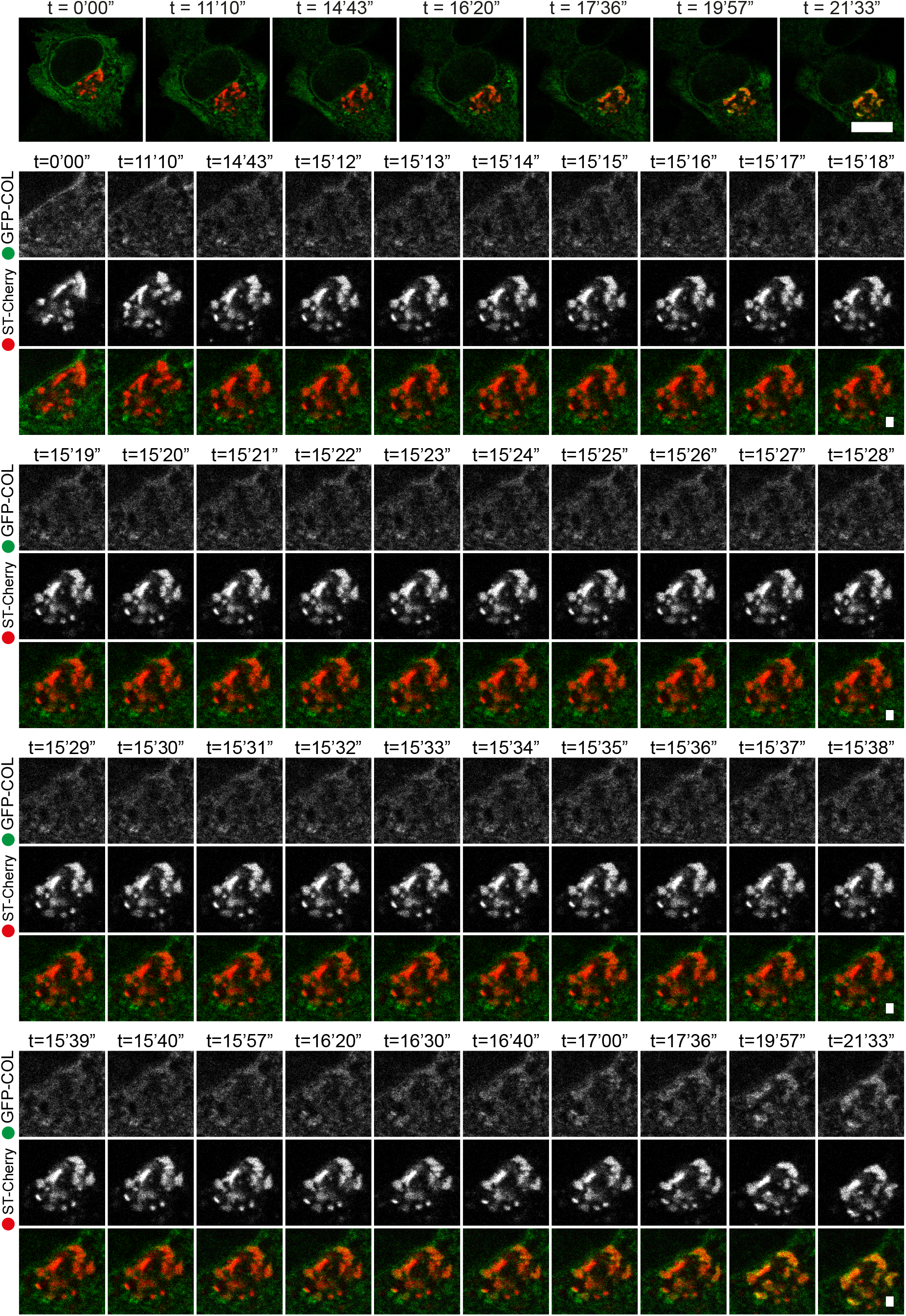
Transport of GFP-COL to the Golgi occurs without the use of large carriers Image stills from confocal live cell imaging of RPE-1 stably expressing GFP-COL (green; GFP-COL-RPE) 18.5 hours post-co-transfection with the trans-Golgi marker ST-Cherry (red; utilising the RUSH system). Timepoints indicate mins after addition of asc/biotin. Whole cell imaged at 1 frame every second (derived from Video 5) with corresponding enlargements with channels for GFP-COL and ST-Cherry in greyscale, as well as the overlay image below. Accumulation and filling of the Golgi with GFP-COL occurs without visible large structures in the proximity of the Golgi. A total of n = 3 was acquired. Scale bars = 10 μm and 1 μm (in enlargements).

**Supplemental Figure S3:**
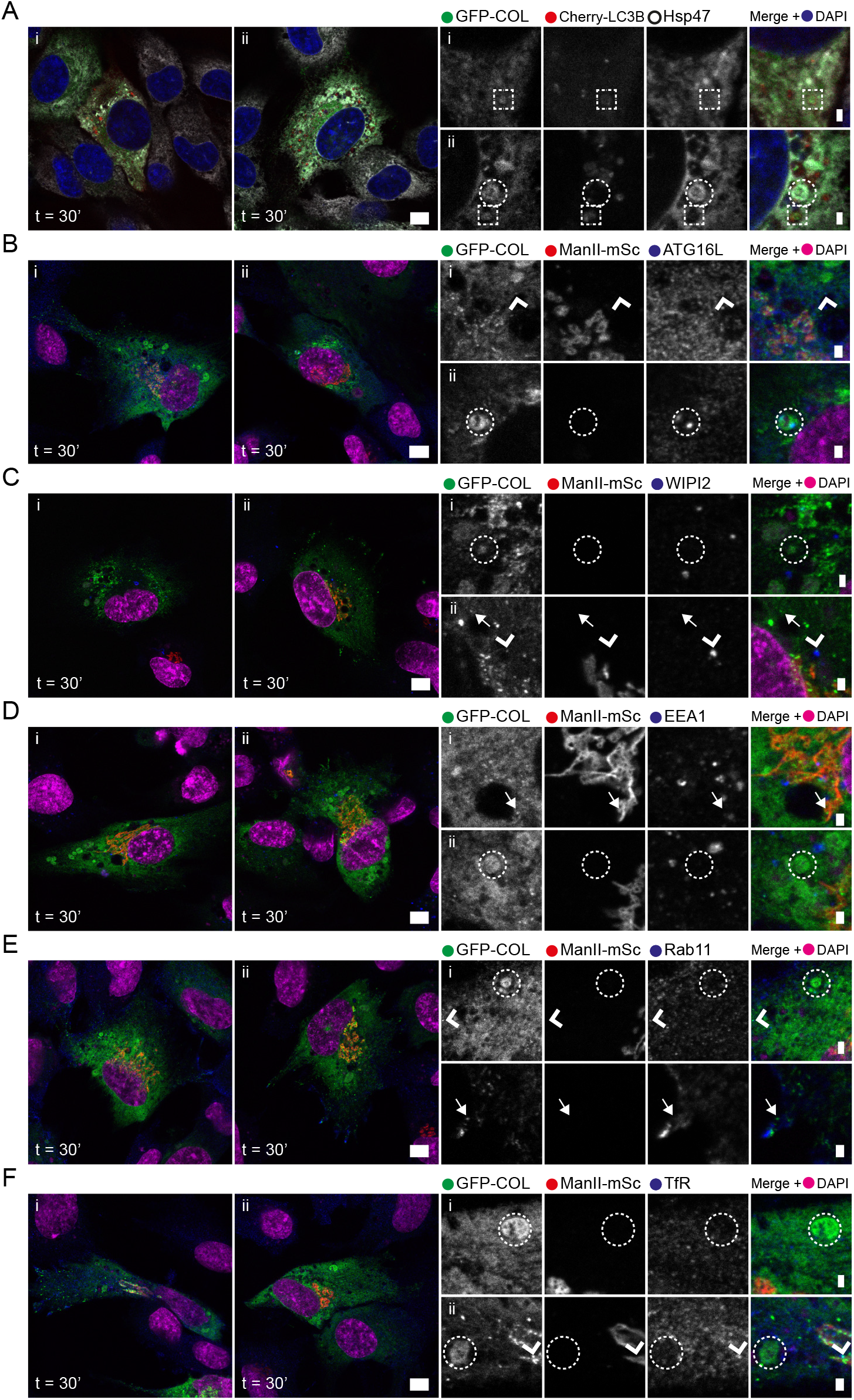
Large GFP-COL structures are not positive for autophagosomal or endosomal markers. Confocal images of GFP-COL-RPE stably expressing GFP-COL (green) co-transfected with either the autophagosomal marker Cherry-LC3B (A) or MannII-mSc (B-F; red) 20- and 18-hours post-transfection, respectively, and labelled for antibodies against autophagosomal and endosomal markers. Cells were fixed with PFA (except for C, which was fixed with methanol) after incubation with asc/biotin (500 μg.ml-1 and 400 μM, respectively) for 30 mins, or just ascorbate when transfected with Cherry-LC3B. Subsequently antibody labelling for Hsp47 is displayed in white, while autophagosomal markers ATG16L and WIPI2 or endosomal markers EEA1, Rab11 or transferring receptor (TfR) are depicted in blue. Panels show two examples of whole cells with corresponding enlargements showing the individual channels as greyscale, followed by the merge image including nuclear labelling with DAPI (imaged as separate channel in pseudo-colour blue (A) or magenta (B – F)). Scale bars = 10 μm and 1 μm for enlargements. Circles indicate large GFP-positive structures not co-localising with autophagosomal or endosomal markers. A: Boxes indicate larger GFP-positive structures co-localising with Cherry-LC3B. These appear negative for Hsp47. B - F: Arrow heads indicate small GFP-positive structures co-labelling for ATG16L, Rab11 or TfR, while arrows highlight small GFP-positive puncta not co-labelling for autophagosomal or endosomal markers, while arrow heads indicate possible colocalising puncta.

**Supplemental Figure S4:**
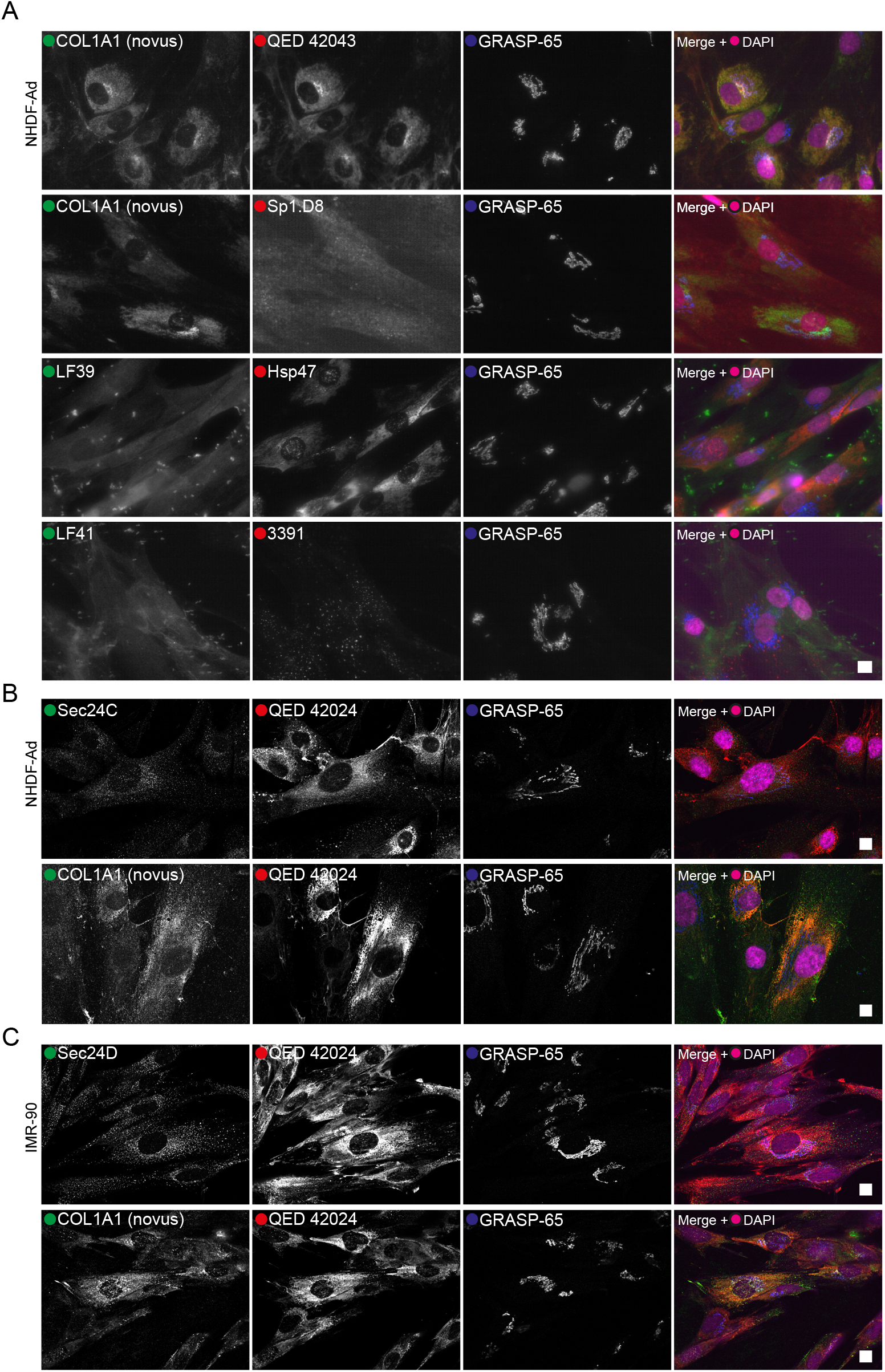
Labelling of collagen type I in fibroblasts using various antibodies shows no apparent large structures. Images of fibroblasts labelled with different antibodies against endogenous procollagen 1α1 / COL1A1, COPII or Hsp47 and cis-Golgi marker GRASP-65 in blue after 30 mins incubation with 500 μg.ml-1 ascorbate prior to fixation with PFA. Rabbit polyclonal antibodies are displayed in green and mouse monoclonal antibodies in red. Panels show the individual channels in greyscale, followed by the overlay image including nuclear DAPI labelling (imaged as separate channel in pseudo colour magenta). Scale bars = 10 μm. A: Widefield images of primary skin fibroblasts NHDF-Ad labelled with COL1A1 antibody (Novus), LF39 and LF41 against the N- and C-terminal propeptide of COL1A1 in green, co-labelled with antibodies against either COL1A1 C-peptide (PIP) (QED 42043), N-telopeptide COL1A1 (Sp1.D8), collagen-specific chaperone Hsp47 or C-terminal COL1A1 (3391). B: Confocal images of NHDF-Ad labelled with a rabbit-polyclonal against the inner layer COPII component Sec24C or COL1A1 (Novus) and mouse monoclonal QED 42024, raised against the C-peptide (PIP) of COL1A1. C: IMR-90 near primary lung fibroblasts labelled with either COL1A1 (Novus) or COPII antibody against Sec24D in combination with labelling against mouse monoclonal QED 42024. B and C were brightness and contrast enhanced using ImageJ. None of the antibody labelling shows apparent large structures labelling for COL1A1 and or COPII.

**Supplemental Figure S5:**
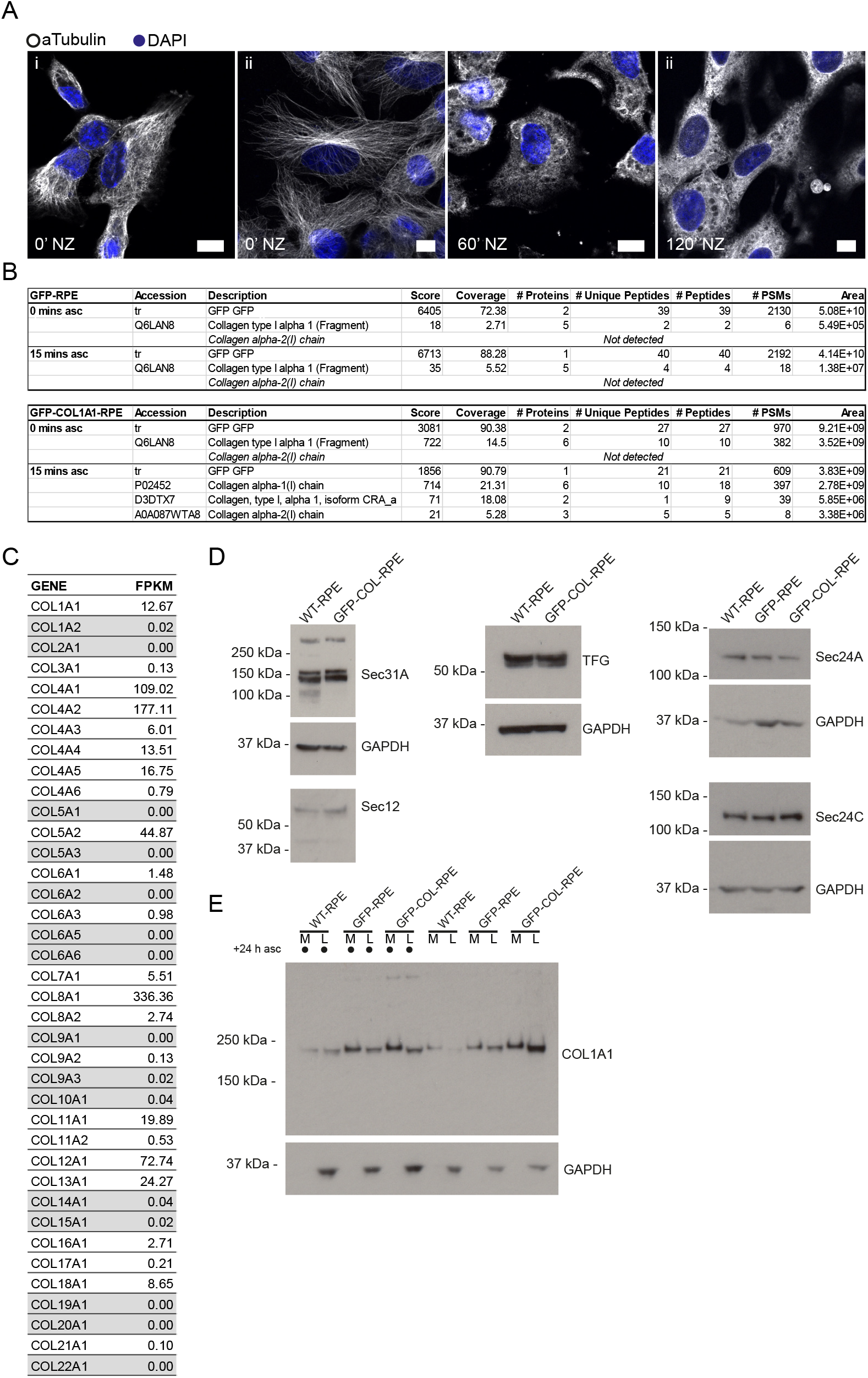
Nocodazole efficacy and collagen and COPII levels in RPE. A: Confocal images of RPE-1 stably expressing GFP-COL (GFP-COL-RPE) 18.5 hours post-co-transfection with the trans-Golgi marker ST-Cherry (both not shown) and labelled with an antibody against α-tubulin (greyscale) and nuclear DAPI staining in blue. Timepoints indicate minutes after addition of nocodazole (NZ) prior to fixation with methanol. The channel for α-tubulin was brightness and contrast enhanced using ImageJ’s autocorrection. A total of n = 3 was acquired. Scale bar = 10 μm. B: Results of a GFP-pull down experiment from GFP-RPE (used as a control) and GFP-COL-RPE incubated without or in presence of 500 μg.ml-1 ascorbate for 15 mins post-24-hour ascorbate flush prior to cell lysis. Columns display the accession number, protein description, score, coverage, the number of proteins, unique peptides and peptides, as well as the number of PSMs and the peak area sorted by number of PSMs. Listed are only the proteins of interest: GFP, different forms of detected COL1A1, as well as COL1A2. COL1A2 could only be detected in the GFP-COL-RPE sample incubated with ascorbate for 15 mins. C: RNA-Seq data from RPE-1 cells listing collagen isotypes detected and the number of fragments per kilobase of exon per million reads mapped (FPKM). Genes with an FPKM below 0.1 are highlighted ingrey. D: Immunoblots of RPE-1 (WT-RPE), GFP-COL-RPE and GFP-RPE cells showing expression levels of components of the COPII machinery Sec31A, Sec12, TFG, Sec24A and Sec24C from cell lysates obtained after 4 days of growth on a 10cm dish and in presence of ascorbate for 24 hours in case of Sec24A and C. GAPDH was used as a loading. E: Immunoblot of cell lysates (L) and media fractions (M) of RPE-1 cells (WT-RPE), GFP-COL-RPE and GFP-RPE cells incubated with or without 50 μg.ml-1 ascorbate for 24 hours before collecting the samples with GAPDH as loading control and COL1A1 (Novus Biologicals) as protein of interest.

## Legends to supplemental material

The following videos correspond to the images shown in figures 1 – 4 and 7, S1 and S2 as mentioned and show trafficking of GFP-COL to the Golgi with addition of ascorbate or asc/biotin (500 μg.ml^-1^ ascorbate and 400 μM biotin, respectively; as does Video 5 for Figure S2 at a higher frame rate), as well as the influence of nocodazole on this process.

### Video 1: Ascorbate- and biotin-dependent trafficking of GFP-COL

Confocal live cell imaging of RPE-1 stably expressing GFP-COL (green; GFP-COL-RPE) 21 hours post-transfection with the *trans*-Golgi marker ST-Cherry (magenta; utilising the RUSH system). Image stills are shown in Fig. 1D. Acquisition at 1 frame every 30 seconds over a period of 49 mins with a playback rate of 12 frames per second. Time stamp shows time after addition of asc/biotin with a final concentration of 500 μg.ml^-1^ ascorbate and 400 μM biotin. A total of n=3 sets was acquired. Scale bar indicates 10 μm.

### Video 2: Transport of GFP-COL to the Golgi using the RUSH system is biotin controllable

Confocal live cell imaging of RPE-1 stably expressing GFP-COL (green; GFP-COL-RPE) 17 hours post-co-transfection with the *cis*-Golgi marker MannII-mSc (magenta; utilising the RUSH system). Time stamps indicate the total imaging time, followed by the time after addition of ascorbate, followed by the time after addition of biotin. **Part 1** shows the cell after addition of ascorbate over a time of 23 mins, with no visible transport to the Golgi. **Part 2** shows the same cell after addition of biotin with a final concentration of 500 μg.ml^-1^ ascorbate and 400 μM biotin, respectively. Accumulation and filling of the Golgi with GFP-COL occurs without visible large structures in the proximity of the Golgi and can be observed at about 5 – 10 mins after addition of biotin. A total of n = 2 was acquired. Acquisition at 1 frame every 18 seconds over a period of about 1 hour with a playback rate of 12 frames per second. Scale bar = 10 μm. Corresponding image stills are shown in Fig. S1A.

### Video 3: Transport of GFP-COL to the Golgi is not cell type specific

Confocal live cell imaging of BJ-5ta transiently expressing GFP-COL (green) and *trans*-Golgi marker ST-Cherry (magenta; utilising the RUSH system) 17 hours post-co-transfection. Time stamp indicates the time after addition of asc/biotin. Accumulation and filling of the Golgi with GFP-COL can be observed at about 10 mins after addition of asc/biotin. Some GFP-positive structures can be identified with an estimated maximum diameter of 400 nm (measurement from the object translocating over the nucleus at about 17 mins post-asc/biotin using ImageJ) that translocate to the Golgi. A total of n = 1 was acquired. Acquisition at 1 frame every 15 seconds over a period of about 1 hour with a playback rate of 12 frames per second. Scale bar = 10 μm. Corresponding image stills are shown in Fig. S1B and images of fixed cells after trafficking in Fig. 3Aii.

### Video 4: Transport of GFP-COL to the Golgi in IMR-90 human lung fibroblasts

Confocal live cell imaging of IMR-90 transiently expressing GFP-COL (greyscale) 20 hours post-transfection. Time stamp indicates the time after addition of ascorbate. Accumulation and filling of the estimated Golgi area with GFP-COL occurs at about 1 hour after addition of ascorbate. A total of n = 1 was acquired. Acquisition at 1 frame every 2.5 seconds over a period of about 1 hour with a playback rate of 12 frames per second. Scale bar = 10 μm. Corresponding image stills are shown in Fig. S1C.

### Video 5: Transport of GFP-COL to the Golgi occurs without the use of large carriers

Confocal live cell imaging of RPE-1 stably expressing GFP-COL (green; GFP-COL-RPE) 18.5 hours post-co-transfection with the *trans*-Golgi marker ST-Cherry (magenta; utilising the RUSH system). Accumulation and filling of the Golgi with GFP-COL occurs without visible large structures in the proximity of the Golgi. Small GFP-positive puncta can be observed traversing in the direction towards the Golgi at about 10 mins asc/biotin (highlighted by arrow heads). A total of n = 3 was acquired. Acquisition at 1 frame every second over a period of 30 mins with a playback rate of 12 frames per second. Time stamp shows time after addition of asc/biotin with a final concentration of 500 μg.ml^-1^ ascorbate and 400 μM biotin and a scale bar of 1 μm. Corresponding image stills are shown in Fig. S2.

### Video 6: GFP-COL transport to the Golgi in RPE1 cells

Confocal live cell imaging of RPE-1 stably expressing GFP-COL (green; GFP-COL-RPE) 23 hours post-transfection with the *trans*-Golgi marker ST-Cherry (magenta; utilising the RUSH system). Image stills are shown in Fig. 2A - B. Cells fixed after live imaging and labelled for giantin are shown in Fig. 3Aii. Acquisition at 1 frame every 20 seconds over a period of 52 mins with a playback rate of 6 frames per second. Time stamp shows time after addition of asc/biotin with a final concentration of 500 μg.ml^-1^ ascorbate and 400 μM biotin. A total of n = 3 sets was acquired. Scale bar = 10 μm.

### Video 7: GFP-COL transport to the Golgi in RPE1 cells

Confocal live cell imaging of RPE-1 stably expressing GFP-COL (green; GFP-COL-RPE) 18 hours post-transfection with the *trans*-Golgi marker ST-Cherry (magenta; utilising the RUSH system). Image stills are shown in Fig. 2C - D. Acquisition at 1 frame every 30 seconds over a period of 19 mins with a display rate of 6 frames per second. Time stamp shows time after addition of asc/biotin with a final concentration of 500 μg.ml^-1^ ascorbate and 400 μM biotin. A total of n = 3 sets was acquired. Scale bar = 10 μm.

### Video 8: GFP-COL transport from the ER to the Golgi, prior to fixation and co-labelling with antibodies of interest

Confocal live cell imaging of RPE-1 stably expressing GFP-COL (green; GFP-COL-RPE) 18 – 24 hours post-transfection with the *trans*-Golgi marker ST-Cherry (magenta; utilising the RUSH system). Time stamps show the time after addition of asc/biotin. Scale bar = 10 μm. A total of n = 3 sets was acquired. The display rate is 6 frames per second. Video 8.i was acquired at 1 frame every 30 seconds over a period of 12 mins (images of cells after fixation are shown in Fig. 3Ai). Video 8.ii was acquired at 1 frame every 40 seconds over a period of 45 mins (images of cells after fixation are shown in Fig. 3B). Video 8.iii was acquired at 1 frame every 30 seconds over a period of 8 mins (images of cells after fixation are shown in Fig. 3Di). Video 8.iv was acquired at 1 frame every 15 seconds over a period of 17 mins (images of cells after fixation are shown in Fig. 3Dii).

### Video 9: Transport of GFP-COL to the Golgi is COPII dependent

Confocal live cell imaging of RPE-1 stably expressing GFP-COL (green; GFP-COL-RPE) 20 hours post-co-transfection with the inner-layer COPII marker mSc-Sec23A (magenta; utilising the RUSH system). Time stamp indicates the time after addition of asc/biotin. Acquisition at 1 frame every 2.79 seconds over a period of about 13.5 mins with a playback rate of 12 frames per second. Scale bar = 10 μm (0.25 μm for the enlargement marked by the square and displayed in the top right corner). A total number of n = 9 was acquired. Accumulation and filling of the expected Golgi area with GFP-COL can be observed at about 2 mins after addition of asc/biotin. The arrow head in the enlargement highlights a GFP-COL structure colocalising with mSc-Sec23A close to the Golgi region at about 1 mins 38 sec before the GFP-COL structure becomes part of the Golgi network at 2 - 3 mins. Corresponding image stills are shown in Fig. 4A - B.

### Video 10: GFP-COL transport to the Golgi is independent of microtubules

Confocal live cell imaging of RPE-1 stably expressing GFP-COL (green; GFP-COL-RPE) 18 hours post-transfection with the *trans*-Golgi marker ST-Cherry (magenta; utilising the RUSH system). Image stills are shown in Fig. 7A - B. Acquisition at 1 frame every 25 seconds over a period of 90 mins with a display rate of 3 frames per second. Time stamp shows time after addition of asc/biotin with a final concentration of 500 μg.ml^-1^ ascorbate and 400 μM biotin and incubation with 5 μM nocodazole for 60 mins. For each set of live imaging experiment 3 cells from the same dish were imaged simultaneously. A total of n=3 sets was acquired. Scale bar indicates 10 μm.

